# Massively Parallel Profiling of RNA-targeting CRISPR-Cas13d

**DOI:** 10.1101/2023.03.27.534188

**Authors:** Hung-Che Kuo, Joshua Prupes, Chia-Wei Chou, Ilya J. Finkelstein

## Abstract

Type VI CRISPR enzymes cleave target RNAs and are widely used for gene regulation, RNA tracking, and diagnostics. However, a systematic understanding of their RNA binding specificity and cleavage activation is lacking. Here, we describe <u>RNA</u> <u>c</u>hip-<u>h</u>ybridized <u>a</u>ssociation-<u>m</u>apping <u>p</u>latform (RNA-CHAMP), a massively parallel platform that repurposes next-generation DNA sequencing chips to measure the binding affinity for over 10,000 RNA targets containing structural perturbations, mismatches, insertions, and deletions relative to the CRISPR RNA (crRNA). Deep profiling of Cas13d, a compact and widely used RNA nuclease, reveals that it does not require a protospacer flanking sequence (PFS) but is exquisitely sensitive to secondary structure within the target RNA. Cas13d binding is strongly penalized by mismatches, insertions, and deletions in the distal crRNA-target RNA regions, while alterations in the proximal region inhibit nuclease activity without affecting binding. A biophysical model built from these data reveals that target recognition begins at the distal end of unstructured target RNAs and proceeds to the proximal end. Using this model, we designed a series of partially mismatched guide RNAs that modulate nuclease activity to detect single nucleotide polymorphisms (SNPs) in circulating SARS-CoV-2 variants. This work describes the key determinants of RNA targeting by a type VI CRISPR enzyme to improve CRISPR diagnostics and *in vivo* RNA editing. More broadly, RNA-CHAMP provides a quantitative platform for systematically measuring protein-RNA interactions.

## INTRODUCTION

Class 2 CRISPR-Cas systems are useful for genetic engineering because they target DNA and/or RNA with a single effector protein^1^. Among class 2 enzymes, Cas13 subtypes exclusively target and cleave RNA^2–9^. Cas13s process their own CRISPR-RNAs (crRNAs), bind a target RNA that is complementary to the crRNA, and cleave the target RNA (*cis*-cleavage) and other RNA molecules via a non-specific RNase activity (*trans*-cleavage)^10,11^. These RNase activities are catalyzed by two Cas13-encoded higher eukaryotes and prokaryotes nucleotide-binding (HEPN) domains which can be mutagenically inactivated to convert Cas13 into an RNA-binding module^10,12–14^. Due to these activities, Cas13 variants are broadly used *in vitro* and in cells^5,15–17^. For example, Cas13d—one of the most compact and biochemically active Cas13 enzymes—can efficiently knockdown RNA in mammalian cells and animal models^5,18–25^. Moreover, Cas13d fusions are used for RNA tracking, editing, modification, and splicing regulation^5,7,26,27^. Cas13d has also been applied for nucleic acid detection in CRISPR diagnostics^28^. However, the binding and cleavage specificity of Cas13d on partially matched target RNAs has not been fully characterized, limiting our understanding and biotechnological applications of this enzyme.

Biochemical studies have reported various targeting specificities across Cas13-family enzymes. Some enzymes require a protospacer flanking sequence (PFS)—a specific sequence adjacent to the target—for RNA cleavage. For example, LshCas13a prefers a non-G 3’-PFS, whereas BzCas13b favors non-G 5’-PFS and 3’PFS of NNA or NAN^2,3,6^. However, LwaCas13a, PspCas13b, and RfxCas13d (CasRx) may not require any PFS at all^2,4,5,7,13,16,17^. The cleavage activity of LwaCas13a, LshCas13a, and LbuCas13a is sensitive to mismatches in the central region of crRNA-target RNA duplex^2,16,29,30^. A large-scale Cas13d screen in mammalian cells also concluded that a distal spacer region (positions 15-21) is largely intolerant to mismatches^31^. These experiments primarily use Cas13 cleavage as an output, conflating binding, activation, and cleavage into a single reporter. Interpreting studies across different experimental conditions and target RNAs is especially challenging because RNA structure can change drastically even with a single nucleotide substitution and may also impact both binding and cleavage. A complete understanding of off-target activity requires the biochemical separation of binding and cleavage across a defined set of structural target RNA and sequence perturbations.

Here, we describe RNA-CHAMP (<u>c</u>hip-<u>h</u>ybridized <u>a</u>ssociation-<u>m</u>apping <u>p</u>latform) for massively parallel profiling of RNA-protein interactions on upcycled next-generation DNA sequencing chips. Using RNA-CHAMP, we characterize how target RNA alterations impact the RNA binding by Cas13d. Contrary to other Cas13-family enzymes, Cas13d does not have a strong PFS preference. However, nucleotide substitutions that increase the overall target RNA secondary structure profoundly decrease the binding affinity. Mismatches and intramolecular basepairing in the distal region of the target RNA strongly decrease Cas13d binding. Surprisingly, mismatches in the proximal region of the target do not affect binding but inhibit nuclease activity. A series of biophysical models of increasing complexity shed insights into the mechanism of Cas13d binding. Together, our results and model suggest that Cas13d initially recognizes the target RNA in the solvent-exposed distal spacer region, followed by RNA duplex formation towards the target RNA in the proximal region. Structural elements in the distal segment impede Cas13d binding. Using these insights, we design a series of partially mismatched crRNAs to detect single nucleotide polymorphisms (SNPs) in circulating SARS-CoV-2 variants. These results will guide future RNA editing and CRISPR diagnostics applications. More broadly, RNA-CHAMP will enable high-throughput mapping of protein-RNA interactions in diverse cellular processes.

## RESULTS

### RNA-CHAMP measures protein-RNA interactions on sequenced Illumina chips

RNA-CHAMP repurposes Illumina next-generation sequencing (NGS) chips to quantify millions of protein-RNA interactions (**Fig. 1A**). RNA molecules are transcribed *in situ* from a template DNA library that has been sequenced using an Illumina MiSeq instrument. We designed the DNA library with the T7 RNA polymerase (RNAP) promoter, a variable region of interest, and the RNAP-stalling *terB* DNA sequence^32,33^. This DNA sequence is recognized by Tus, a bacterial protein that blocks T7 RNAP translocation^34,35^. The identity and physical coordinates of each DNA cluster are determined during next-generation sequencing (NGS). After sequencing, the chip is regenerated to remove leftover fluorescent nucleotides and resynthesize the double-stranded (ds) DNA^36^. Tus is then added to the chip to stall T7 RNAP. *In vitro* transcription and subsequent stalling of T7 RNAP tethers the transcript to its DNA template.

**Figure 1.**
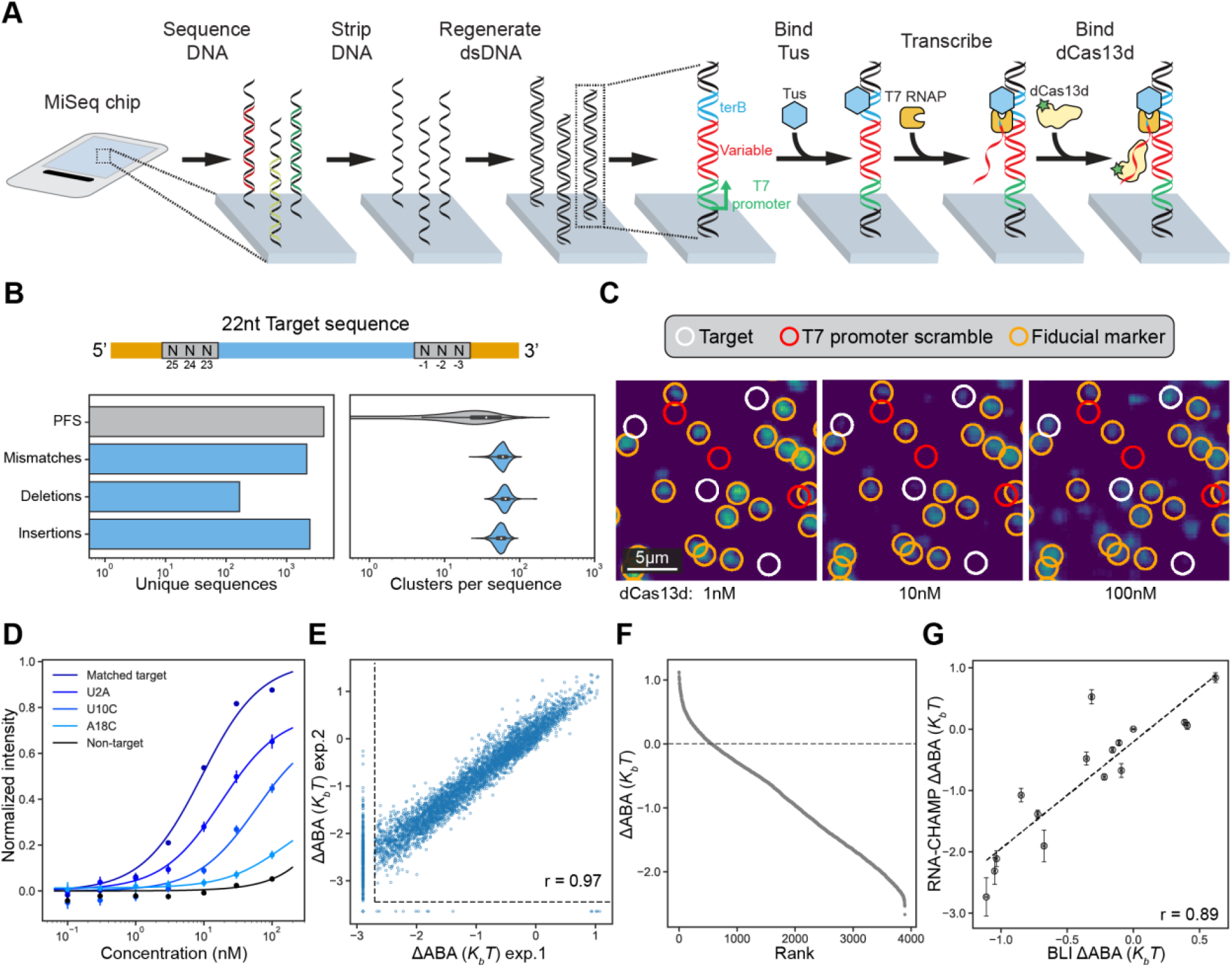
Massively parallel protein-RNA profiling via RNA-CHAMP. **(A)** RNA-CHAMP workflow. DNA is regenerated on the surface of a sequenced MiSeq chip and is transcribed with T7 RNA polymerase (RNAP). Tus retains T7 RNAP and the associated transcript on the DNA. Fluorescent Cas13d is incubated in the chip and the chip surface is imaged. **(B)** Top: Schematic of the RNA library. Spacer sequences (orange) include Illumina primer annealing sites and the T7 RNAP promoter. The 22-nucleotide target RNA (blue) is flanked on both ends by three random nucleotides (PFS, gray). Bottom: Summary of the unique DNA sequences in the synthetic library (right), and the number of clusters observed via NGS for each unique library member (left). **(C)** Fluorescent images of the chip surface after incubating with increasing Cas13d concentrations. White circles: library clusters. Red circles: scrambled promoters that cannot produce RNA. Orange circles: fiducial markers used for image alignment. **(D)** Quantification of fluorescent intensities for the indicated mismatch sequences. For example, U2A indicates a U to A substitution at the second position in the target RNA. Solid lines are fit to a hyperbolic function. Data are shown as median ± S.D. from all cluster intensities (n > 10). **(E)** Correlation of two independent RNA-CHAMP experiments. Dashed lines denote the limit of detection. Pearson’s *r* = 0.97. **(F)** Rank-ordered graph of the ΔABA for ~4,000 library members. The dashed line represents the ΔABA of the matched target (MT). Sequences below our detection limit are omitted. **(G)** Correlation of the ΔABA and biolayer interferometry (BLI)-determined binding affinities. Error bars are the standard deviation of ΔABA (RNA-CHAMP) and fit 95% confidence interval (BLI) (>10 clusters for RNA-CHAMP, three concentrations for BLI). The dashed line is the linear fit of data points. Pearson’s *r* = 0.89.

We first assayed the efficiency of RNA stalling in MiSeq chips. To confirm that Tus recognizes *terB*-encoding DNA clusters, we purified FLAG-epitope labeled Tus and fluorescently labeled it with ATTO488-conjugated anti-Flag antibody^37^ (**Fig. S1A**). We sequenced a library that included DNAs with and without the *terB* sequence. Over 90% of *terB*-encoding DNA clusters co-localized with fluorescent Tus (**Fig. S1A**). The remaining *terB*-encoding clusters could not be resolved by our image processing software, usually due to the spatial overlap of two or more clusters. Importantly, Tus did not bind clusters that lacked *terB*. All downstream analysis was conducted on *terB*-containing DNA clusters. To confirm that transcripts are stably retained after in vitro transcription (IVT), they were hybridized with a complementary ATTO647N-labeled oligonucleotide *in situ* (**Fig. S1B**). The chip also included DNA clusters with scrambled T7 RNAP promoters as negative controls. We observed an RNA signal from ~90% of promoter-containing clusters, but not from scrambled promoter clusters (**Fig. S1B**). These results demonstrate that RNA-CHAMP can generate libraries of user-defined RNA molecules on repurposed MiSeq chips.

Next, we characterized the specificity and off-target RNA binding of *Eubacterium siraeum* (*Es*) Cas13d, a prototypical member of the CRISPR RNA-guided RNA nucleases^4,5^. We purified nuclease-dead *Es*Cas13d with an N-terminal SNAP-tag and fluorescently labeled it with SNAP-Surface-488 (hereafter referred to as “dCas13d”; **Fig. S1C**). The ribonucleoprotein (RNP) complex was reconstituted to 100% homogeneity by incubating dCas13d with a 4-fold excess of the crRNA followed by size exclusion chromatography. Native gel electrophoresis confirmed complete RNP formation (**Fig. S1D**). This procedure was repeated for RNPs with different crRNAs and used in all subsequent experiments. The SNAP-tag did not alter the protein’s RNA-binding affinity, as measured via Biolayer Interferometry (BLI) (**Fig. S1E**).

Type VI CRISPR-Cas nucleases recognize a protospacer-flanking sequence (PFS) that is immediately adjacent to the 5’ or 3’ of the target RNA^2–7^. To test whether *Es*Cas13d is sensitive to the PFS, we included three randomized bases on both the 5’ and 3’ of the matched target sequence. In addition, the target RNA library included up to two mismatches, insertions, or deletions relative to the crRNA (**Fig. 1B & Supplemental File 1**). To confirm that our findings are generalizable across targets, we also prepared a second library with a different target RNA sequence but identical design characteristics (**Figs. S3 & S4**). We sequenced both RNA libraries to ensure >10 DNA clusters for all library members (**Fig. 1B, right**). We also included unrelated DNA sequences as controls or fiducial markers for downstream image analysis and spatial registration. After sequencing, the MiSeq chip was regenerated and transcribed with T7 RNAP for downstream experiments.

Transcribed libraries were incubated with increasing concentrations of dCas13d (**Figs. 1C, D**). Clusters with T7 promoters showed dCas13d concentration-dependent increases in fluorescence intensities, whereas scrambled promoters showed no dCas13d binding (**Fig. 1C**). The fluorescent intensities of clusters across all concentrations were background-subtracted and fit with a Hill equation without cooperativity to determine the apparent binding affinity (ABA) (**Fig. 1D** and **Methods**)^36,38^. To directly compare the relative binding affinity across the entire library, we calculated the change in the binding affinity (ΔABA) as the natural logarithm of the matched target affinity divided by partially matched RNA library members (see **Methods**). Two biological replicates showed excellent reproducibility across the entire dynamic range of binding affinities (**Fig. 1E**). In partially matched libraries, we measured the binding affinities for 3,893 sequences from the target library out of 4,936 total members (**Fig. 1F**). The remaining target RNA sequences had binding affinities or fluorescent signals that were below our detection limit. Using BLI, we validated a subset of 16 RNA targets across the entire dynamic range of the RNA-CHAMP experiments, including sequences with mutations in the target RNA as well as the PFS (**Fig. S2**). ABAs calculated from BLI measurements were in excellent agreement with the sequences from our library, indicating that RNA-CHAMP accurately captures the relative affinities of dCas13d to its target RNA sequences (Pearson’s *r* = 0.89; **Fig. 1G, S2**). Moreover, the BLI analysis indicates that the ΔABA is dominated by *k_on_*, likely because the target RNA-crRNA duplex is extremely stable after hybridization (**Table S1**). We conclude that RNA-CHAMP is a quantitative platform for massively parallel protein-RNA interaction profiling.

### Cas13d requires a partially unstructured target RNA

We measured dCas13d binding affinity with a PFS library consisting of three random nucleotides on the 5’ and 3’ end of the 22 nt matched target sequence (target #1) (**Fig. 2**). We measured the ΔABA for 1457 PFS combinations. The remaining sequences were below our detection threshold. Although dCas13d exhibited a ~3-fold difference in ΔABAs across the entire PFS dataset, it did not have a strong PFS preference (**Fig. 2A**). We observed a similar result in a second target (target #2) library but with a slight preference for non-G 3’-PFS at position 1 (**Fig. S3**). Combining the top 25% highest ABA binding sequences in both targets confirm that Cas13d has a weak preference for the 3’-PFS (**Fig. S3D**). This preference, however, only partially explains the wide range of ΔABAs for these datasets.

**Figure 2.**
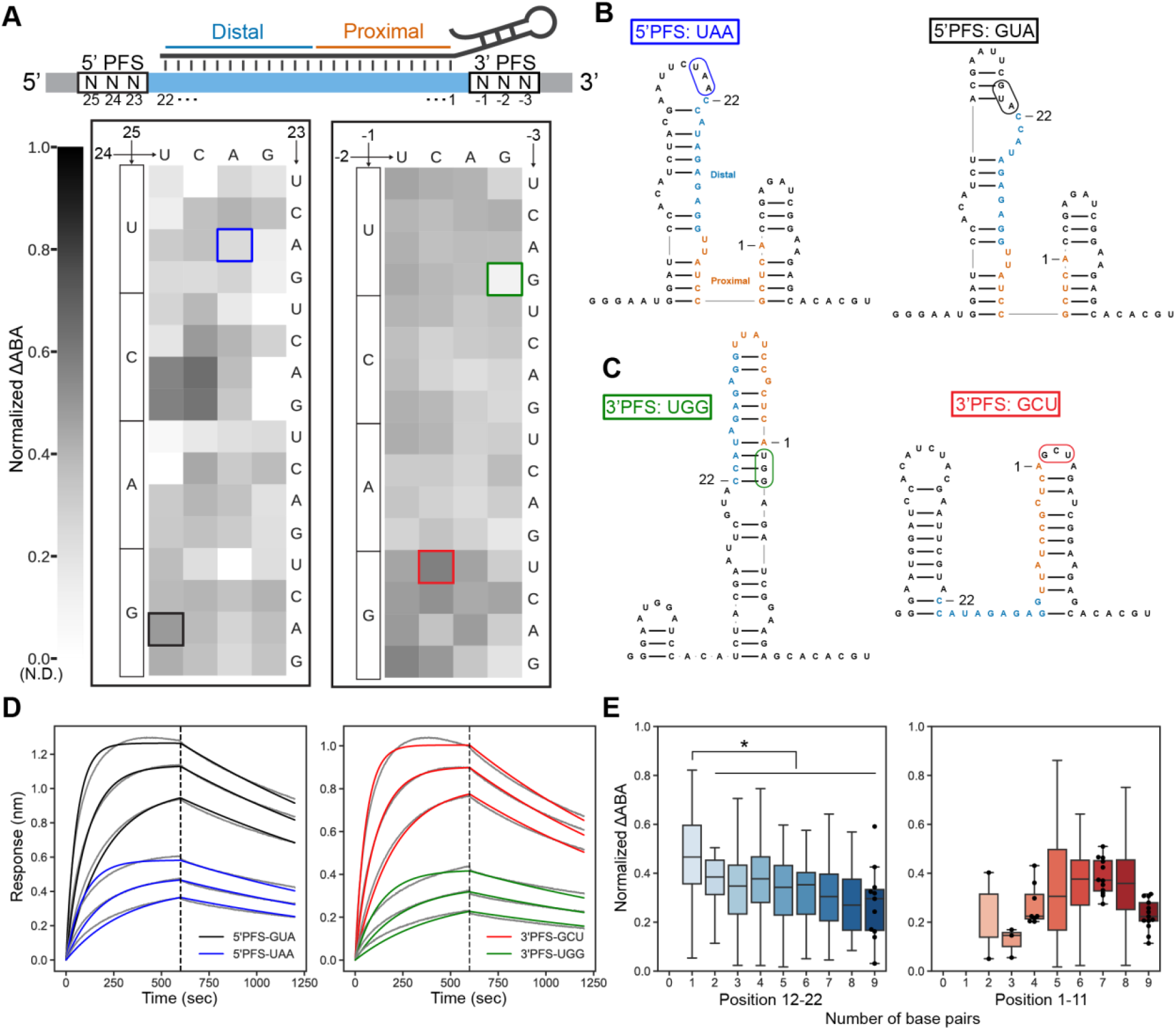
Target RNA structure is a strong determinant of Cas13d binding affinity. **(A)** Top: Schematic of the target RNA library. The target RNA is perfectly matched to the crRNA. Bottom: Normalized ΔABAs for the 5’ PFS and 3’ PFS. In the plot, each block in the heat map is the mean of all detectable sequences with that 5’ PFS (left) and 3’ PFS (right). All sequences were normalized to the scale of zero to 1 for easy comparison between targets. **(B)** Secondary structure predictions of two illustrative examples for 5’ PFS. Left: a low-affinity target RNA (5’-UAA). Right: a high-affinity target RNA (5’-GUA). The PFSs are boxed in blue and black in (A). **(C)** Predicted secondary structure of a low-affinity PFS (3’-UGG; left), and a high-affinity PFS (3’-GCU; right). The PFSs are boxed in green and red in (A). **(D)** BLI curves of the highlighted PFS sequences in (B) and (C). Grey lines are experimental curves. Colored lines are the global fit to a 1:1 binding model. **(E)** Normalized ΔABA of PFS sequences grouped by their base pairing count within the target region. Graph of the base pairing count in positions 12-22 (Left) and 1-11(Right). Error bars are the standard deviation of normalized ΔABA. Swarm plots are used when the number of sequences is less than 20.

We reasoned that the target RNA secondary structure can regulate Cas13d binding^29,31,39^. When inspecting both high- and low-affinity target RNAs, we observed that dCas13d prefers target RNAs that are not predicted to be basepaired in the distal regions (positions 11-22) (**Figs. 2A-C & S3A, B**)^40^. For example, the 5’-PFS GUA forms a stem with the 5’ constant region and exposed positions 19-22, which resulted in ~2-fold stronger dCas13d binding than 5’-PFS UAA (**Fig. 2A, B**). Similarly, 3’-PFS GCU forms a stem with the 5’ constant region and exposed position 14-20. These exposed distal nucleotides in 3’-PFS GCU led to a ~2-fold increase in dCas13d binding affinity relative to 3’-PFS UGG (**Fig. 2A, C**). BLI measurements independently validated these observations (**Fig. 2D**). This also confirms that low-affinity PFSs have a similar off-rate (*k_d_*), but slower on-rates (*k_a_*) than high-affinity PFSs (**Fig. 2D, Table S1**). These results highlight that RNA structure regulates Cas13d access to the matched target RNA.

To determine how the local target RNA structure affects Cas13d binding, we computed the number of predicted basepairs in its proximal (positions 1-11) and distal (positions 12-22) regions. Basepairs can form with the RNA outside the target, or within the target itself. For example, the 3’-GCU PFS sequence in **Figure 2C** has three proximal and eight distal basepairs, whereas the 3’-UGG PFS has nine proximal and six distal basepairs. Increased basepairing in the distal region of the target RNA decreased the ΔABA. In contrast, no statically significant trend was identified between the number of basepairs and the ΔABA in the proximal region (**Fig. 2E, S3C**). Cas13d prefers to engage the distal end of the target RNA first; this region must remain partially unstructured for efficient binding (see **Discussion**). Taken together, we conclude that Cas13d does not have a PFS requirement but prefers to bind target RNAs with unpaired distal nucleotides.

### Cas13d is sensitive to mismatches in the distal region of the target RNA

To determine how Cas13d binds off-target RNAs that resemble the target sequence, we constructed a library comprised of 66 single mismatches, 2079 double mismatches, and 2439 insertions & deletions relative to the crRNA within the 22-nt target sequence (target #1) (see **Fig. 1B**). For all experiments, the 5’- and 3’-PFS remained constant. Of the 4936 library members, we measured ΔABAs for 3893 target RNAs. 1043 sequences didn’t significantly change the dCas13d fluorescent signal, even at the highest RNP concentrations. **Figure 3A** summarizes two biological replicates of the ΔABA for all possible single mismatches. We also measured ΔABAs across a similarly designed library but with different target crRNA sequences (target #3) (**Fig. S4**). The binding trends were broadly the same across these two libraries.

**Figure 3.**
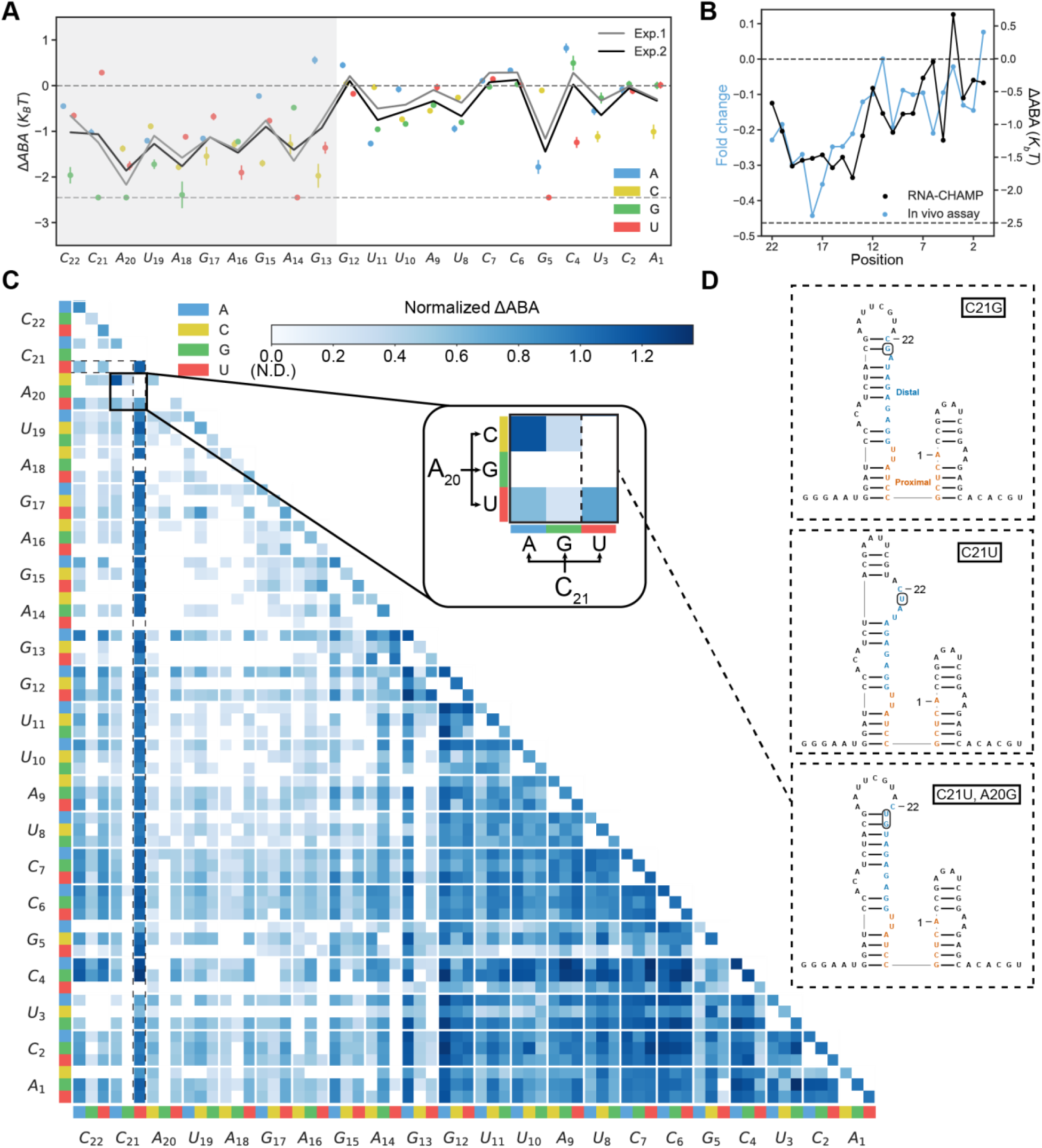
Cas13d binding is sensitive to mismatches in the 5’-region of the target RNA. **(A)** Summary of single mismatch-dependent changes in the ΔABA for two biological replicates. Upper dashed line: matched target ΔABA, lower dashed line: RNA-CHAMP detection limit. **(B)** Comparison of ΔABA and *in vivo* cleavage of a reporter gene (data adapted from 31). For RNA-CHAMP, all three possible mismatches were averaged at each position along the target RNA. **(C)** Normalized ΔABAs of all double mismatched sequences normalized to the matched target. Inset: blowup of all possible mismatches at target positions A20 & C21. **(D)** Secondary structure predictions of three illustrative examples. Top: C21G. Middle: C21U. Bottom: C21U, A20G. The mismatches are boxed.

We first analyzed the impact of a single mismatch between the target and crRNA (**Fig. 3A**). Mismatches at positions 13-22 significantly decreased the ΔABA. In contrast, single mismatches at positions 1-12 had little to no effect on binding compared to the matched target (**Fig. 3A**). The identity of the mismatch at the same position led to profoundly different outcomes. For example, a C21U substitution has a similar ΔABA to the matched target, but binding was virtually undetectable with C21G. The C21U substituted is predicted to match the structure of the matched target (**Fig. 3D**, middle). Moreover, C21U creates a G-U wobble basepair with the crRNA, which yields a similar binding affinity to the matched target. C21G, in contrast, creates additional intra-molecular basepairs at positions 19-21 (**Fig. 3D, top**). In a dataset with a different crRNA-target pair, we saw a similar but slightly broader sensitivity region to mismatches at positions 9-20 (**Fig. S4A, B**). We compared our binding results to a dataset of RfxCas13d RNA cleavage activity reported in mammalian cells (**Fig. 3B**)^31^. Because this dataset used different target RNA sequences, we compared the mean ΔABA from all three mismatches across two targets to the mean cleavage activity at each position along the RNA target. RNA knockdown efficiency in mammalian cells is reduced when mismatches are in the distal position, analogously to our binding data (**Fig. 3B**). Overall, Cas13d can tolerate G-U wobble basepairs and shows a strong sensitivity to distal mismatches.

Next, we analyzed the impact of two mismatches on Cas13d binding affinity (**Fig. 3C**). Binding was largely unaffected if both mismatches occurred in positions 1-12 (dark blue squares in **Fig. 3C**). We observed multiple instances where the RNA structure drastically changed the ΔABA. Such sequences appear as “stripes” of strong color in **Fig. 3C**. For example, C21U with an additional substitution (highlighted in dotted line **Fig. 3C**) does not affect the ΔABA compared to the matched target. However, a second mismatch (A20G) in addition to C21U ablates dCas13d binding due to increased basepairing in the distal region of the target RNA (**Fig. 3C, D**). Overall, we observed that Cas13d prefers unpaired distal RNA sequences. We also observed a strong dependence on RNA structure with the second RNA library (target #3). This target RNA is highly folded, with only bases 20-22 not participating in intramolecular basepairing, reducing overall Cas13d affinity (**Fig. S4B, C**). For this RNA structure, some substitutions (e.g., U2A, G4U) relax the proximal to center region of the target RNA (positions 1-13) structure and result in an increased binding affinity relative to the matched target (**Fig. S4C**). Cas13d binary structures suggest that positions 4-8 and 14-20 of the crRNA are solvent exposed and accessible to the environment^13,14^. We speculate that the center exposed region likely contributes to the increased binding affinity. In sum, local RNA structure dominates Cas13d binding affinity. The distal segment of the target RNA must remain partially unpaired for high-affinity binding.

dCas13d retains a high affinity for targets with proximal insertions or deletions (**Figs. S5, S6**). However, insertions and deletions at the distal side of the target RNA on both targets were not tolerated (**Fig. S5, S6)**. We also observed a strong effect from RNA secondary structure. For example, inserting a C between positions 19 and 20 reduces the predicted basepairing at bases 13-20, which increases the ΔABA relative to the matched target (**Fig. S5B**). A G-insertion at the same position leads to undetectably low binding due to new basepairs formed in the distal side of the target RNA (**Fig. S5B**). We observed similar effects of RNA structure on binding affinity in a second RNA target library (target #3) (**Fig. S6A, B**). A C-insertion at position 3 exposes the proximal region that retains similar affinity to the matched target, while a U-insertion increases basepairing and results in undetectable binding. Taken together, these results again show that Cas13d binding is sensitive to distal alterations and local secondary structure. We speculate that Cas13d has a distal seed region and initiates crRNA-target RNA duplexes starting primarily from the distal region (see Discussion).

### Target RNA basepairing is a quantitative predictor for Cas13d binding affinity

We developed a series of linear regression models of increasing complexity to quantitatively understand how mismatches and RNA structure affect Cas13d binding (**Fig. 4A**). Unlike machine learning approaches (also considered below), these models can elucidate the mechanism of Cas13d binding to partially matched targets. The simplest model (Model I) assigns a position-specific penalty for each intramolecular basepair in the predicted target RNA structure (see **Methods**)^40^. This model requires a total of 22 adjustable parameters, one for each nucleotide along the target RNA. In Model II, we add the predicted minimum free energy (MFE) of the entire 73-nt transcript RNA to capture the overall secondary structure. Model III encodes sequence changes relative to the matched target using a relative encoding strategy (see **Methods** and **Fig. S7**). Model IV adds the target RNA’s MFE as another parameter to the relative encoding. Model V combines the basepairing penalty and relative encoding. Finally, Model VI includes all three components: basepairing penalty, relative encoding, and the MFE (**Figs. 4B, C**). We trained each model on half of 4,862 partially matched target sequences across two RNA targets (targets #1 & #3). The resulting model was tested on the withheld half of the sequences in our datasets. After fitting the data, each model’s performance was evaluated by Pearson correlation and information loss via Akaike information criterion (AIC) (**Figs. 4B, C**)^41^.

**Figure 4.**
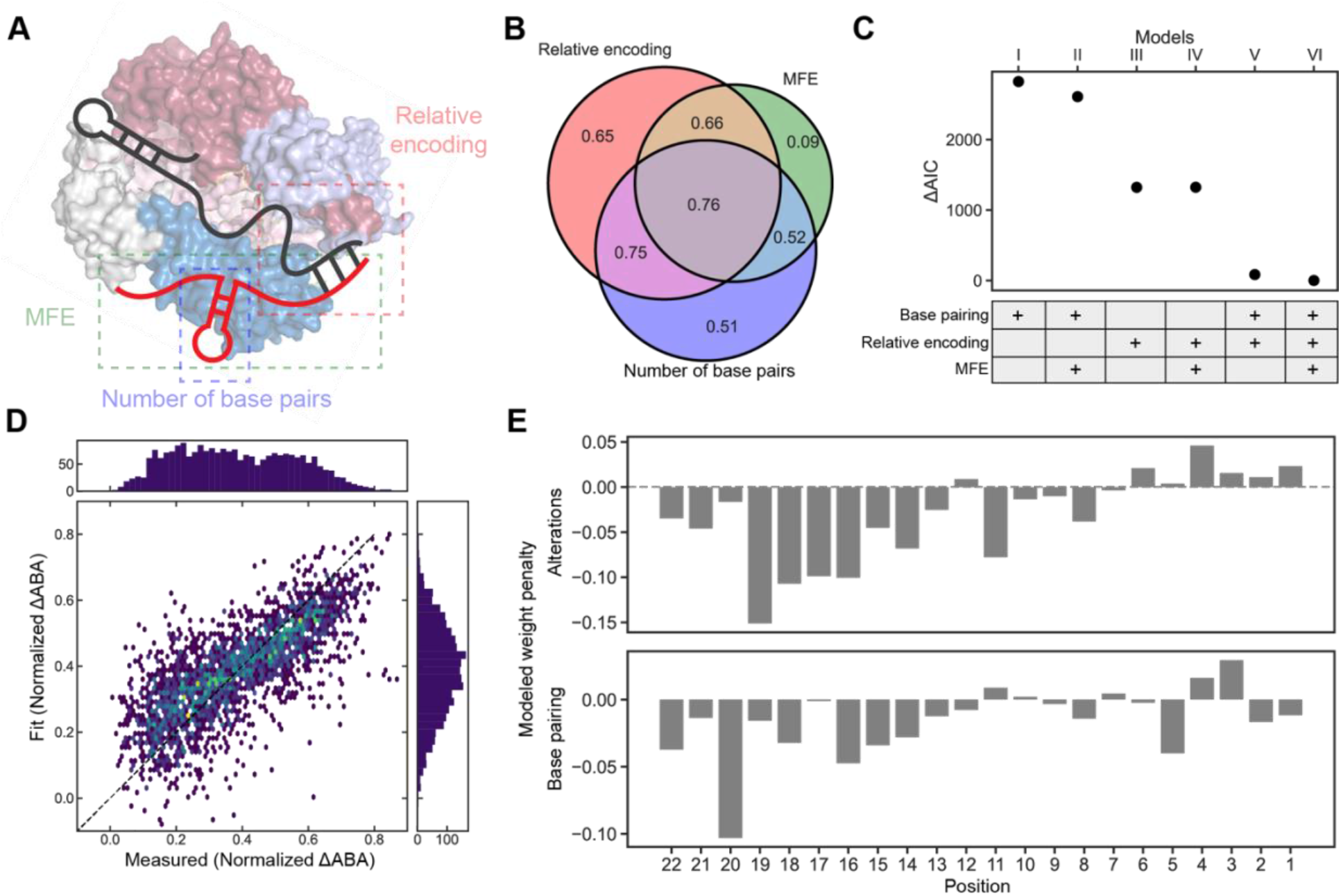
Modeling Cas13d binding. **(A)** Schematic of the three components of our quantitative Cas13d binding models. Relative encoding is the difference between a given sequence and the matched target sequence. The predicted minimum free energy (MFE) of a target RNA is generated by ViennaRNA 2.0^40^. The number of basepairs the counts of the intramolecular RNA base pair in the target region. **(B)** Venn diagram of Pearson’s *r* correlation coefficients from three main components. Correlation between the measured and predicted data are shown in the Venn diagram. **(C)** Akaike information criterion (AIC) of the six models used in this study. ΔAIC is the difference between model I-V and model VI. **(D)** Correlation between the measured and predicted normalized ΔABAs from model VI. Pearson’s *r* = 0.77. **(E)** Weight penalty of all alterations (mismatches, insertions, and deletions) in model VI.

Model I, which only considers intramolecular target RNA basepairing within the 22 nt target sequence result in a Pearson’s *r* = 0.51. Adding the MFE—a measure of the overall structural stability—only weakly improved the correlation and AIC, indicating that local RNA structure is more important than its global stability. Relative encoding has a lower AIC and a Pearson’s *r* = 0.65, performing better than the structure-only model. Finally, combining structural features with relative encoding (Models V & VI) improves both the AIC and Pearson’s *r* to 0.75. Adding the MFE (model VI) slightly improved the AIC, indicating that position-specific mismatches and intramolecular basepairing propensity are sufficient to describe most of the variance in the ABAs. (**Figs. 4B, C**). We also trained a convolutional neural network machine learning (ML) model on the data (**Fig. S7**). Despite having a much larger number of adjustable parameters, the ML model is only marginally better than model VI (Pearson’s *r* = 0.77). Since the ML model’s parameters are not easily interpretable, it doesn’t reveal the mechanisms of RNA binding. Therefore, we dissect Cas13d binding affinities using Model VI below.

We first compared the average penalty for mismatches and indels along the 22 nt target sequence (**Fig. 4E, top**). Cas13d binding is heavily penalized with mismatches or indels at positions 13-22 along the target RNA. In contrast, mismatches at positions 1-12 only minimally decreased the ΔABA. Likewise, basepairing within the target RNA nucleotides 14-22 reduces the ΔABA and is heavily penalized by the model (**Fig. 4E, bottom**). Baspairing within positions 1-13 slightly reduced the ΔABA in the model. Based on these results, we conclude that distal positions 12-22 of the crRNA-target RNA duplex act as an internal “seed” where Cas13d initiates target RNA recognition (see **Discussion**).

### Proximal mismatches suppress Cas13d’s nuclease activity

Next, we tested how mismatches affect Cas13d’s cleavage activity. We measured the cleavage rates of nine mismatched target RNAs that have also been assayed via RNA-CHAMP and BLI (**Fig. 1G**). Time-dependent cleavage of a 6-carboxyfluorescein (6-FAM)-quencher RNA can be followed via an increase in the FAM signal after the fluorophore is released from the quencher (**Fig. 5A**)^28^. The cleavage rate is monitored via the initial slope of the time-dependent fluorescent signal. Cleavage rates were generally correlated with ΔABAs, with three notable outliers. Proximal mismatches C2G, C4A, and C7A did not impact RNA binding but only weakly cleaved the reporter RNA. In contrast, distal mismatches decrease both the binding and cleavage rates (**Fig. 5B-D**). We hypothesize that mismatches at proximal positions disrupt the protein-RNA interface required for activation of the HEPN domain.

**Figure 5.**
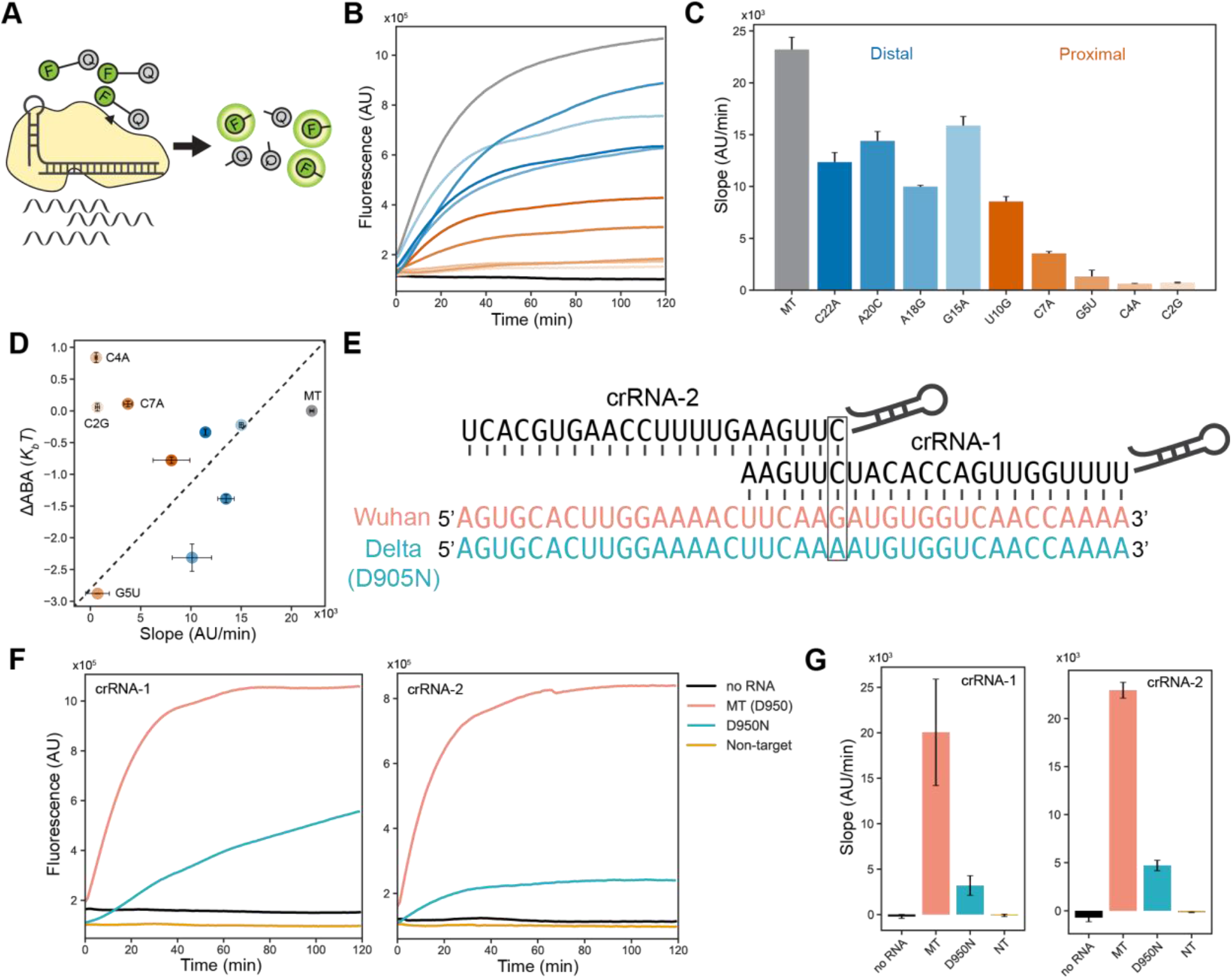
Proximal mismatches limit Cas13d cleavage activity. **(A)** Schematic of the collateral cleavage assay. **(B)** Fluorescent cleavage time courses for matched target and nine mismatched target RNAs. Blue lines are distal mismatched sequences (positions 12-22). Red lines are proximal mismatched sequences (positions 1-11). **(C)** Initial slopes of the traces in **(B)**. Slopes are calculated by the fluorescence changes during the first 20 minutes of the cleavage reaction. Data are shown as mean and S.D. from two replicates. **(D)** Correlation of the cleavage slope with binding affinity (ΔABA). A subset of target RNAs retain strong binding but are cleavage-inactive. Data are shown in mean ±S.D. from two cleavage reactions (x-axis). **(E)** Schematic of mismatch-defined differentiation between SARS-CoV-2 variants of concern (VOC). **(F)** Fluorescent cleavage time courses for SARS-CoV-2 Wuhan and Delta VOCs. **(G)** The initial slope of the trace in **(F)**. Slopes are calculated by the fluorescence changes during the first 20 minutes of the cleavage reaction. Data are shown as mean and S.D. from three replicates.

Cas13d’s mismatch sensitivity can be exploited to rationally design assays that detect single nucleotide polymorphisms (SNPs) in a target RNA. As a proof of principle, we positioned the SNP in the crRNA-target RNA duplex to differentiate between two SARS-CoV-2 variants of concern (VOC) (**Fig. 5E**). Here, the matched target is from the spike gene of the original (“Wuhan”) SARS-CoV-2, but harbors a single G to A substituting (D950N in amino acid sequence) in the position 1 and 17 in the Delta VOC. We designed two crRNAs that position this single nucleotide polymorphism (SNP) within a sensitive region of Cas13d activity, thereby reducing Cas13d cleavage ~5-fold for the Delta VOC RNA (**Figs. 5F, G**)^4,5,28^. This proof-of-principle demonstrates that Cas13d-based diagnostics can be used to distinguish between SNPs by precisely positioning the expected mismatched positions relative to the crRNA.

## DISCUSSION

RNA-CHAMP is a massively parallel platform for probing protein-RNA interactions on used NGS chips. Unlike earlier approaches, CHAMP does not modify any Illumina hardware and is compatible with modern sequencers and chip configurations^32,33,42–44^. Imaging biomolecules on upcycled NGS chips can be adapted by any laboratory with a commercial fluorescence microscope that is capable of either TIR- or epi-illumination and a wide-field camera^36,38,52^. In addition to profiling protein-DNA and protein-RNA interactions, related methods have been adapted for peptide display and other imaging applications^53–55^. We envision that the high optical quality and surface passivation of commercial Illumina chips will extend to massively parallel single-molecule imaging.

Using RNA-CHAMP and quantitative modeling, we show that Cas13d has a “seed region” that prefers a relaxed structure at the distal end of the target RNA (**Fig. 6**). This region is analogous—but not functionally identical—to the PAM-adjacent seed found in Cas9 and DNA-binding CRISPR enzymes^56–60^. The impact of the Cas13d seed is especially profound when the target RNA is perfectly matched with the crRNA. Strong intramolecular basepairing due to changes in the PFS reduces Cas13d binding by over 3-fold relative to a perfectly matched target. Our results highlight that future studies must also consider how the target RNA structure changes enzyme activity. Mismatches can increase the binding affinity when they coincidentally relax intramolecular basepairing within the target. By separating the effect of RNA structure on binding and cleavage, our results explain prior observations that minimal secondary structure in the target RNA correlates with higher cleavage activity in bacterial and mammalian cells^4,29,31,39^.

**Figure 6.**
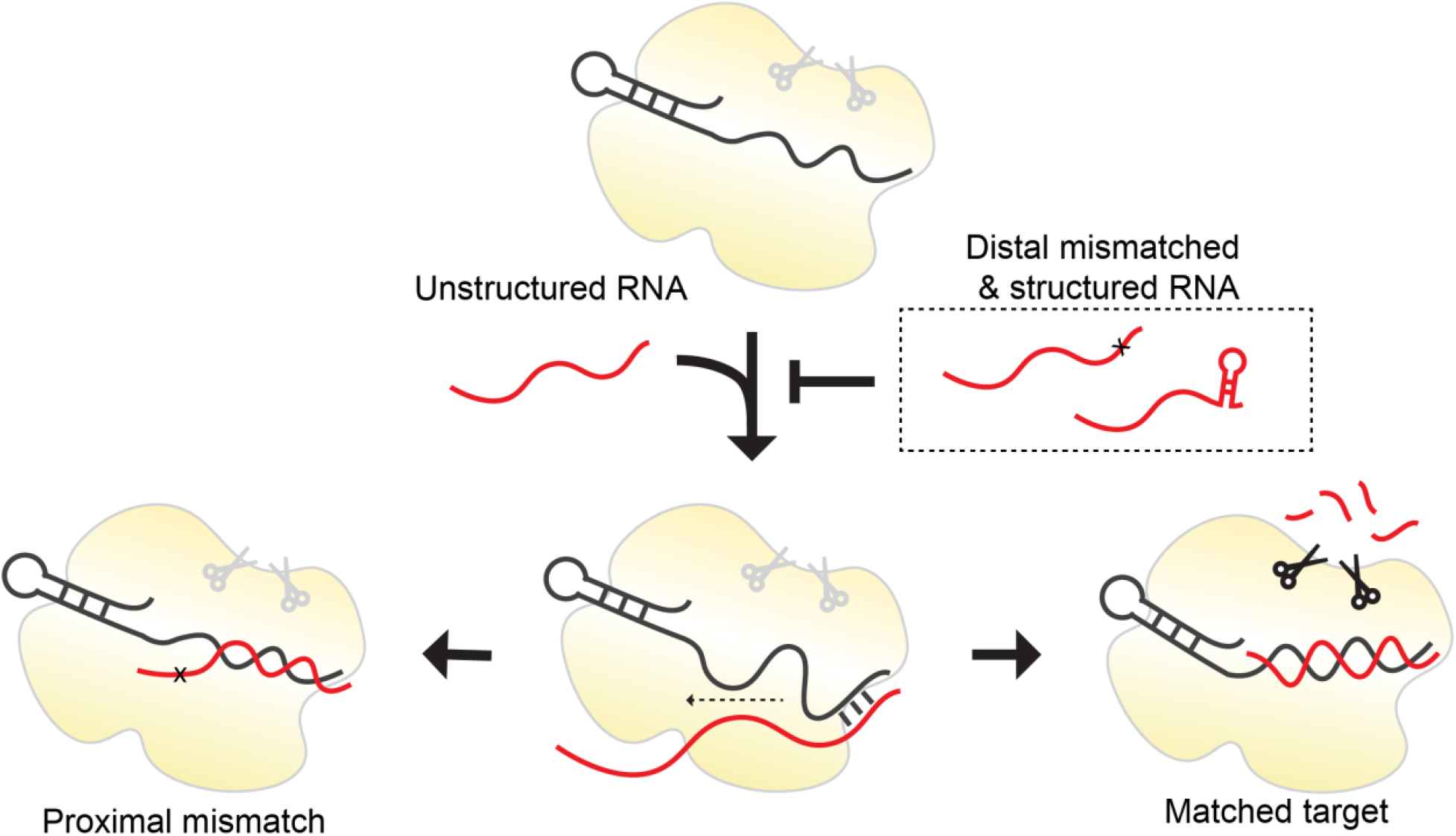
Cas13d binding and nuclease activation follow distinct rules. Cas13d binding is penalized by distal RNA structures and mismatches with the crRNA. After initial distal recognition, the RNA duplex forms outward from the distal position. A mismatch in the proximal region fails to activate the nuclease activity, leading to a catalytically inactive enzyme. Matched target sequences that form a complete RNA duplex activate the nuclease activity.

We separately dissect RNA binding and cleavage to reveal that a subset of mismatched sequences can bind with high affinity but fail to activate the nuclease domain (**Fig. 6**). Cas13d requires basepairing in positions 1-6 to activate its nuclease activity. We leverage this sensitivity to develop guides that can discriminate between circulating SARS-CoV-2 variants. Similarly, LbuCas13a positions 5-8 are critical for cleavage but not binding^30^. This may act as an additional mechanism to suppress nuclease activation and subsequent cell death in prokaryotic hosts. Mismatch-dependent cleavage inactivation may be a universal feature of type VI effectors.

We conclude that Cas13d binding and nuclease activation are governed by distinct spacer-target regions. Mismatches and structural elements in the distal region inhibit binding, whereas proximal mismatches block nuclease activation. These effects, along with the biophysical models developed here, can be selectively used to fine-tune knock-down efficiency in cells by programming mismatches along the crRNA-target RNA duplex. A similar approach has been used to fine-tune CRISPRi with nuclease-dead Cas9 in mammalian cells^61^. In addition, a complete understanding of Cas13d binding and activation can be used for sensitive SNP detection in CRISPR diagnostics (**Fig. 5**)^15^. The structural basis for type VI nuclease activation and the implications for gene editing and prokaryotic immunity are exciting areas for future research.

## SUPPORTING INFORMATION

This article contains Supplemental Figures S1-S7 and Supplemental Tables S1-S3, Supplemental Methods, and a Supplemental Data file. Source code associated with this work is available on GitHub: https://github.com/finkelsteinlab/RNA-CHAMP.

## ACKNOWLEDGMENTS

We thank the staff of the University of Texas at Austin Genomic Sequencing and Analysis Facility, Dr. Rick Russell, and members of the Finkelstein laboratory for carefully reading the manuscript.

## AUTHOR CONTRIBUTIONS

H.-C.K., J.P., C.-W.C., and I.J.F. designed the research. H.-C.K. performed and analyzed the experiments, wrote the bioinformatics software, and performed biophysical modeling. J.P. implemented the biophysical models. C.-W.C. purified proteins and performed biochemical experiments. I.J.F. secured funding. H.-C.K. and I.J.F. wrote the paper with assistance from all co-authors.

