## supplemental material for "Massively Parallel Profiling of RNA-targeting CRISPR-Cas13d"

### SUPPLEMENTARY FIGURES

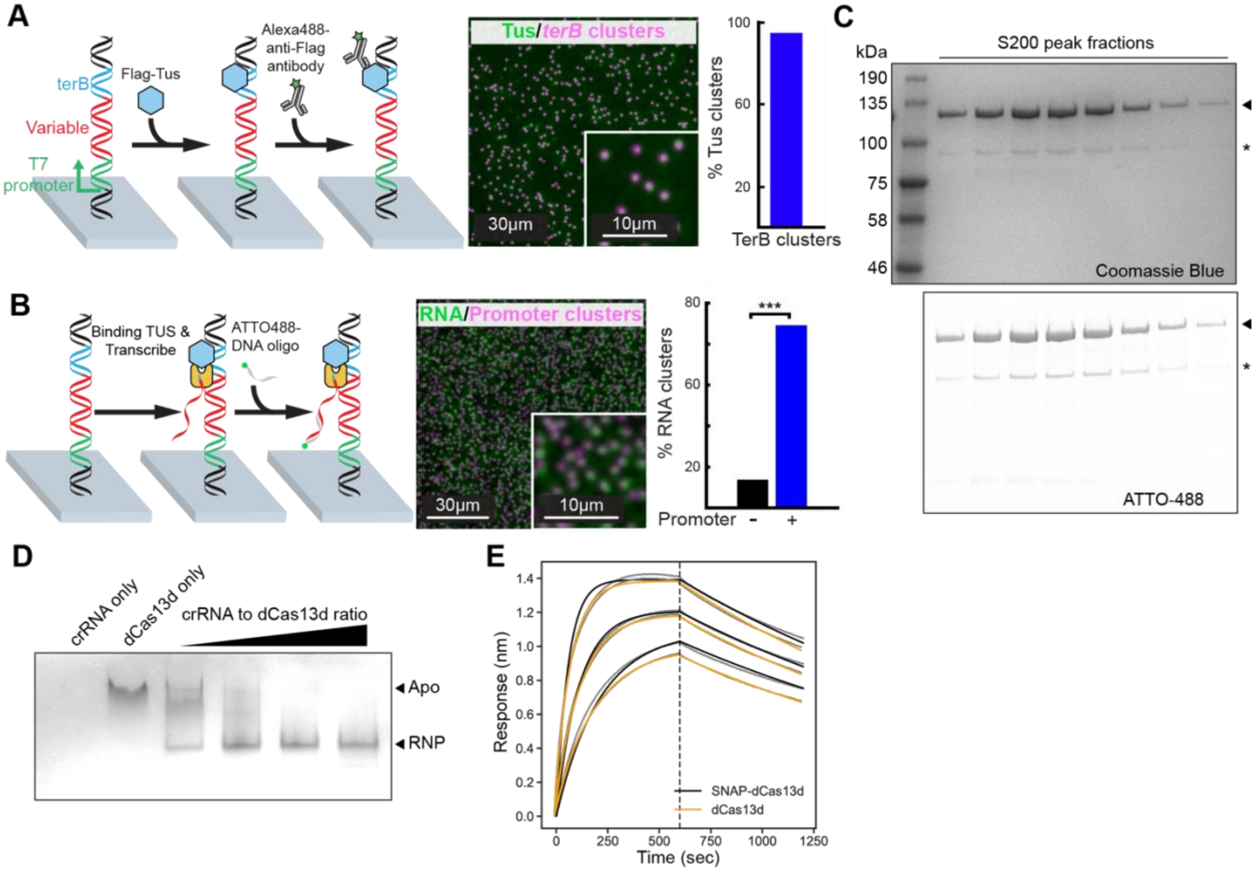


**Figure S1. RNA-CHAMP: A platform for massively parallel measurement of Cas13d binding. (A)** Left: Schematic of the experiment. Middle: Microscope field-of-view. Right: Quantification of fluorescent Tus (green) binding to *terB*-containing DNA clusters (magenta) in a MiSeq flowcell. The bar graph shows the percentage of *terB*-encoding clusters that also had a fluorescent Tus signal. **(B)** Left: Schematic of the experiment, Middle: Microscope field-of-view. Right: Quantification of RNA transcription in a MiSeq flowcell. The bar graph shows the percentage of clusters that hybridized with a ATTO-488 labeled complementary oligo (green) to promoter containing clusters (magenta) and clusters without a T7 promoter. **(C)** Purification of ATTO-488 labeled SNAP-dCas13d. Fractions were collected from an S200 gel filtration column and resolved on an SDS-PAGE gel. Top: Gel visualized by Coomassie blue staining. Bottom: fluorescent image before Coomassie blue staining. Triangle: ATTO-488 labeled SNAP-dCas13d; *: minor truncation product. **(D)** Purification of ATTO-488 labeled dCas13d conjugated with a CRISPR RNA (crRNA). Purified dCas13d was incubated with various amount of crRNA. The crRNA:dCas13d binary complex was separated from apoCas13d via size exclusion chromatography and analyzed in a native tris-glycine gel following by Coomassie blue staining. **(E)** BLI results of SNAP-dCas13d and non-tagged dCas13d. The SNAP tag does not affect Cas13d's binding affinity to RNA.

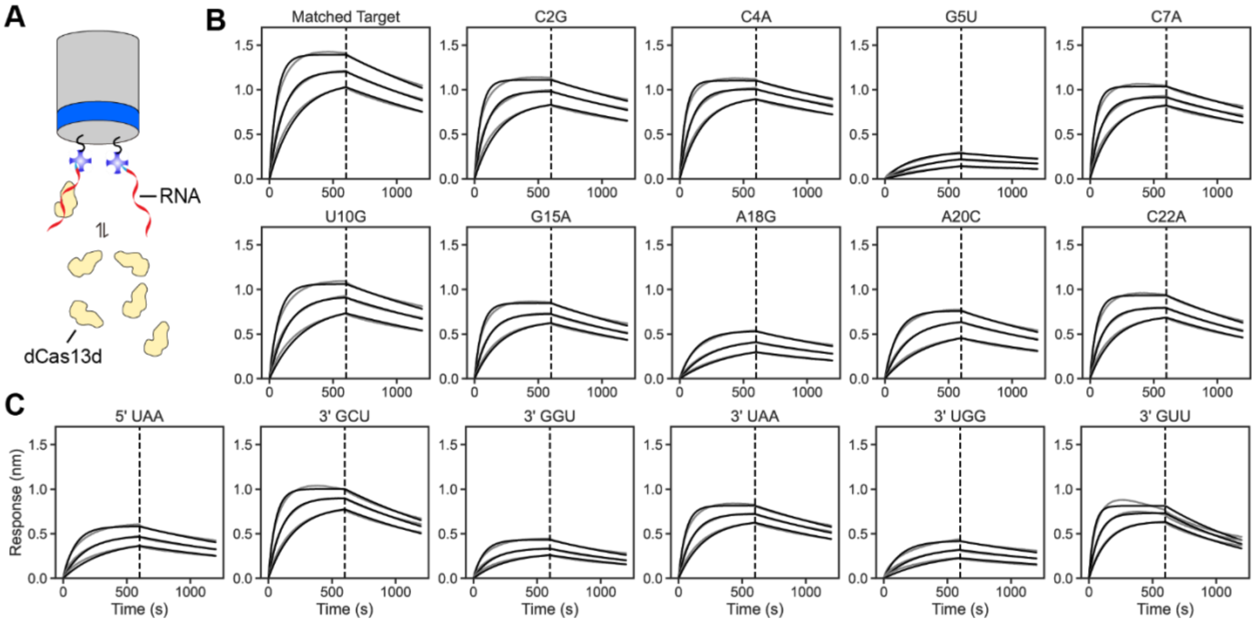


**Figure S2. Biolayer interferometry (BLI) of a subset of the target RNA library.**
**(A)** Schematic of the BLI assay. Biotinylated target RNA is immobilized on a streptavidin-functionalized BLI tip. Separate tips are used for each target RNA. Tips are dipped into a solution containing dCas13d RNPs to monitor the association rate, $k_a$. Dissociation, $k_d$, is monitored by transferring the tip to a buffer solution without any free RNP. For each target sequence, all measurements were repeated with 100 nM, 50 nM, and 25 nM dCas13d. BLI results for **(B)** partially matched and **(C)** PFS sequences. RNA-loaded biosensors were immersed in dCas13d solutions for 600 sec (dashed line). Biosensors were then immersed in the binding buffer for $k_d$ measurements. Black lines are global simultaneous fit to three concentrations. Grey lines are raw BLI curves. Fit results are summarized in Table S1.

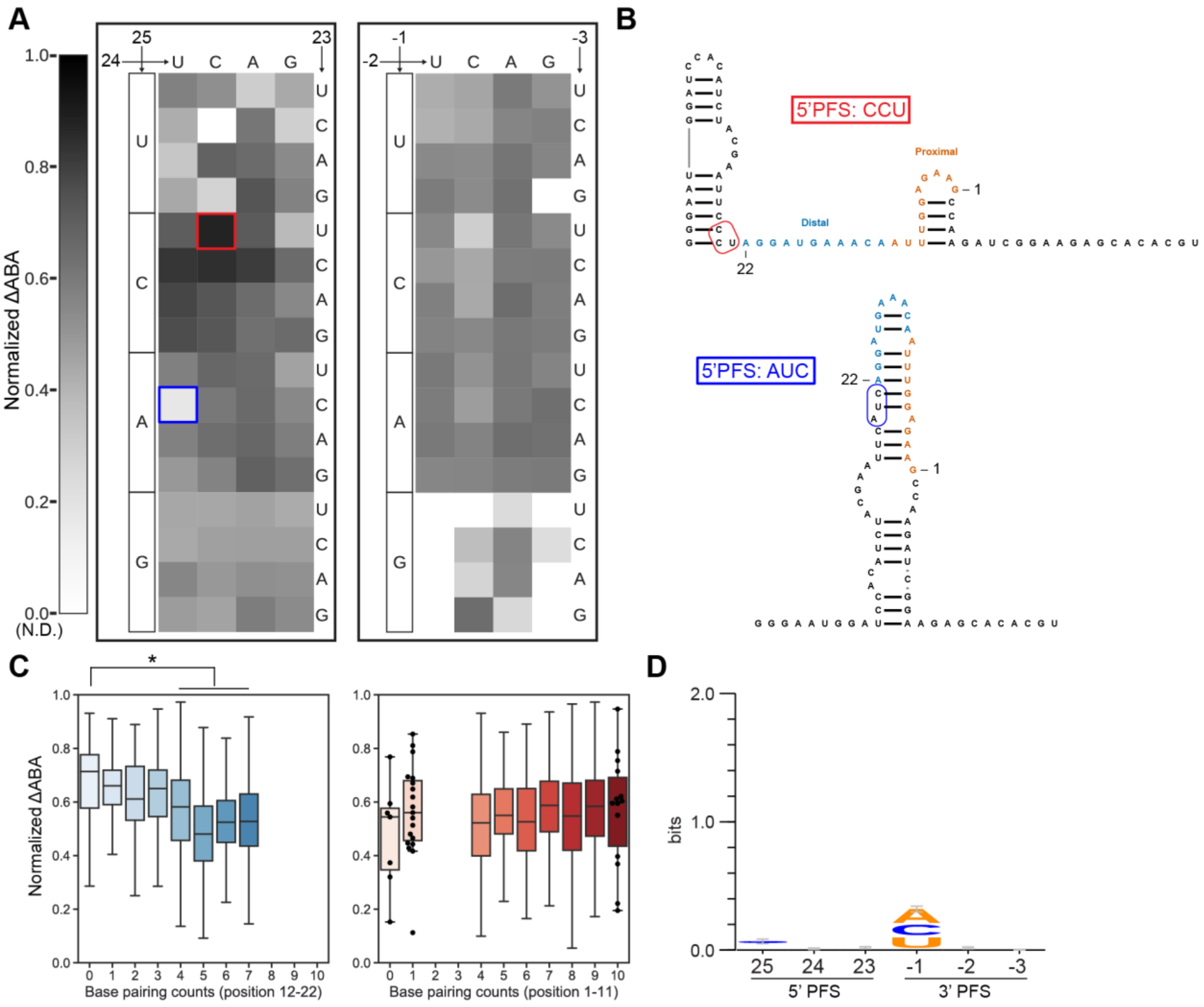


**Figure S3. Cas13d PFS binding analysis on a second target RNA library.**
**(A)** Normalized ΔABAs of Cas13d binding to a PFS library with a second target RNA (left: 5' PFS; right: 3' PFS). Blue and red boxes indicate two targets that are highlighted in the next panel. These sequences have a matched target RNA but significant changes in binding affinities. **(B)** The ViennaRNA-predicted secondary structures of 5'PFS-CCU (red box) and 5'PFS-AUC (blue box). Light blue indicates target RNA sequence. **(C)** Normalized ΔABA of PFS sequences grouped by their basepairing count in the target region. Left: basepairing counts in positions 12-22; right: positions 1-11 of the target RNA. Error bars are the standard deviation of normalized ΔABA. Statistical analysis was performed using unpaired Student's t-test, *$p < 0.05$. **(D)** Sequence logo of the 25% highest affinity PFS sequences across two target RNA libraries from **Figures 2A** & **S3A**.

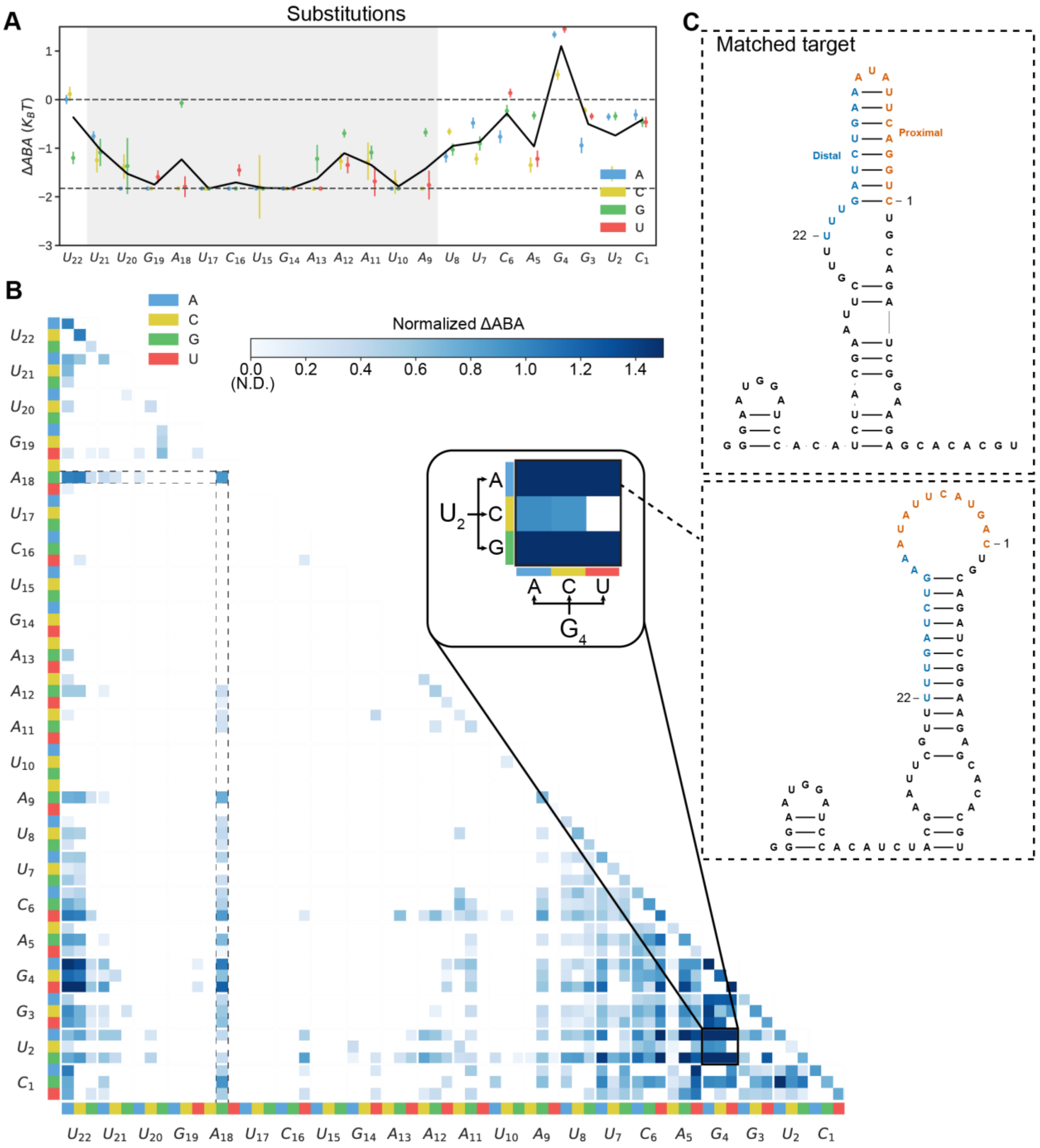


**Figure S4. Mismatch analysis of on a second target RNA library.**

**(A)** Summary of single mismatch-dependent changes in the ΔABA for the second target RNA. Solid black line is the average of all possible substitutions at each position. Upper dashed line is the matched target ΔABA and lower dashed line is the RNA-CHAMP detection limit. **(B)** The normalized ΔABA for all double substitutions (normalized to the matched target). Inset: blowup of all mismatches at target positions $U_2$ & $G_4$. N.D. (not determined) refers to sequences that bound Cas13d with a lower affinity than our detection limit. **(C)** Substitutions that increase intramolecular target RNA basepairing drastically increased the ΔABA. Top: Predicted matched target RNA structure. Bottom: Predicted U2A, G4U structure. Despite two mismatches, this partially matched target binds Cas13d better than the matched target due to relaxed intramolecular basepairing.

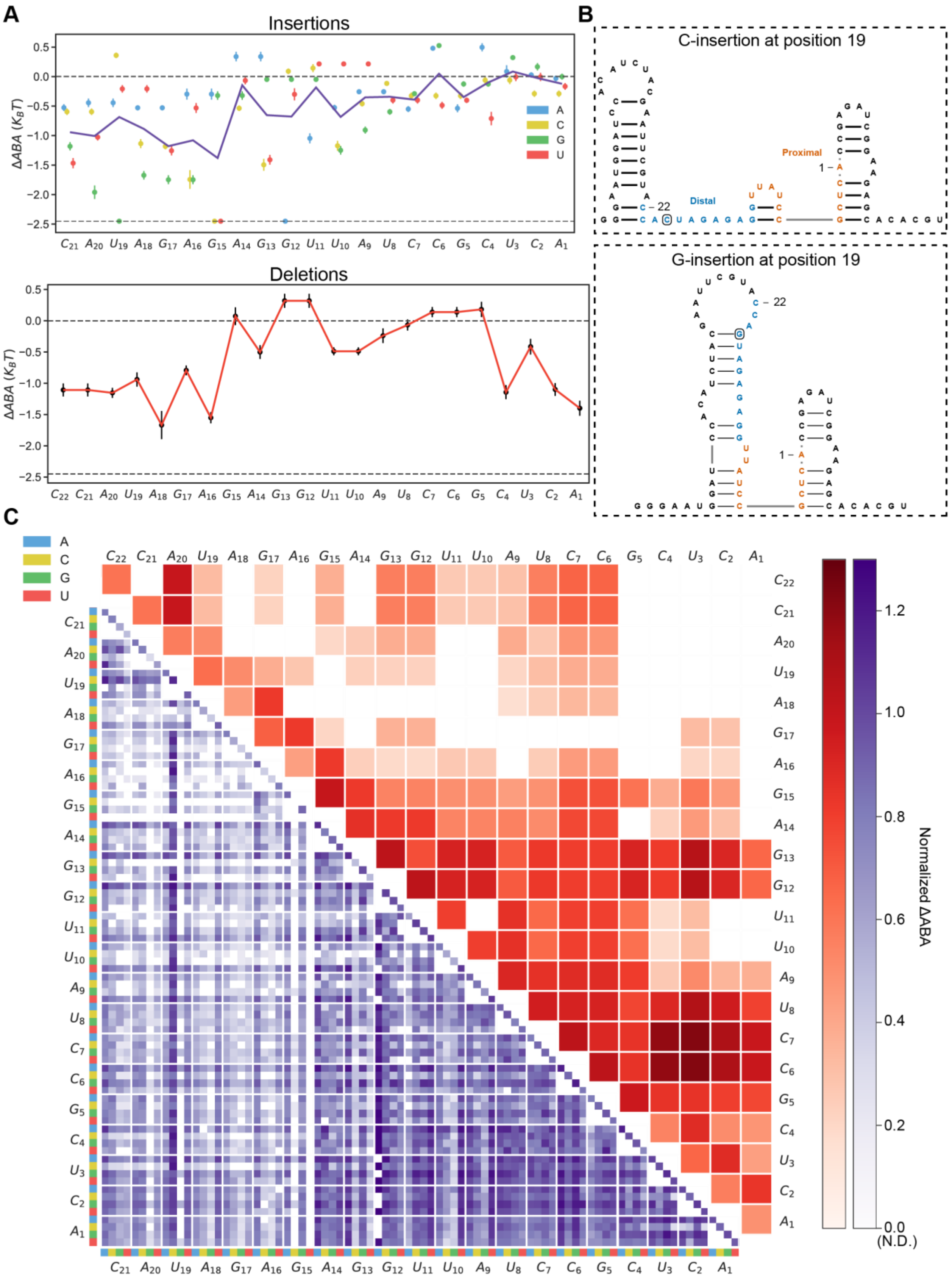
A
Insertions
Deletions
ΔABA ($K_BT$)
B
C-insertion at position 19
G-insertion at position 19
Distal
Proximal
C
Normalized ΔABA
(N.D.)

**Figure S5. Insertion and deletion analysis for the first target.**
**(A)** Changes in the ΔABA for all possible insertion (top) and deletion (bottom). For insertions, the line is an average of the four possible insertions at each position. Upper dashed line is the matched target ΔABA and lower dashed line is the RNA-CHAMP detection limit. **(B)** Predicted structures of two insertions at position 19 that relax (19C, top) or further basepair (19G, bottom) with the target RNA. These changes have a drastic impact on the ΔABA. **(C)** Top triangle plot: double deletion analysis. Bottom triangle plot: double insertion analysis.

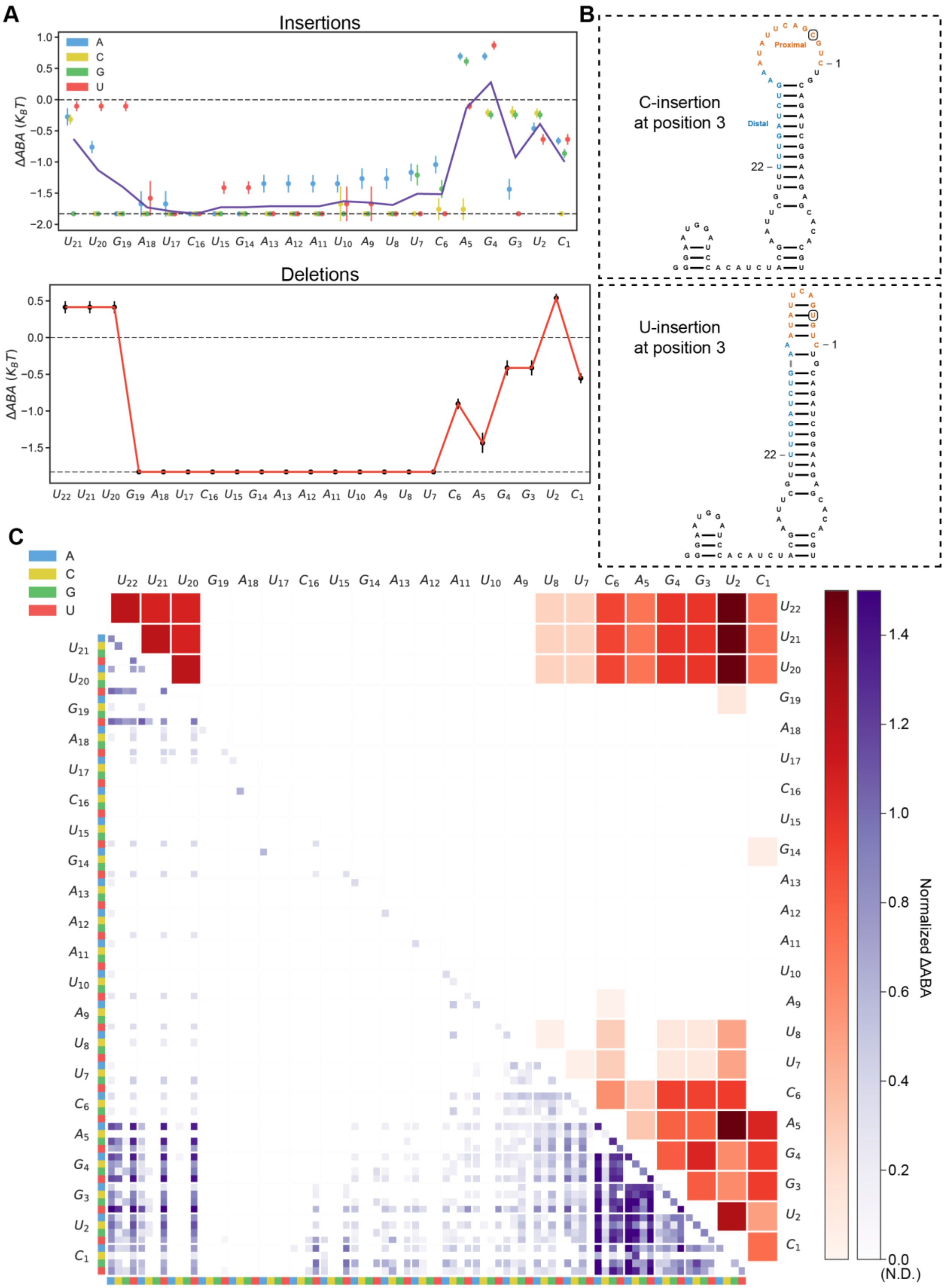

A
Insertions
A
C
G
U
ΔABA (K_BT)
1.0
0.5
0.0
−0.5
−1.0
−1.5
−2.0
U21 U20 G19 A18 U17 C16 U15 G14 A13 A12 A11 U10 A9 U8 U7 C6 A5 G4 G3 U2 C1
Deletions
ΔABA (K_BT)
0.5
0.0
−0.5
−1.0
−1.5
U22 U21 U20 G19 A18 U17 C16 U15 G14 A13 A12 A11 U10 A9 U8 U7 C6 A5 G4 G3 U2 C1
B
C-insertion
at position 3
Proximal
Distal
22 –
– 1
U-insertion
at position 3
22 –
– 1
C
A
C
G
U
U22 U21 U20 G19 A18 U17 C16 U15 G14 A13 A12 A11 U10 A9 U8 U7 C6 A5 G4 G3 U2 C1
U21 U20 G19 A18 U17 C16 U15 G14 A13 A12 A11 U10 A9 U8 U7 C6 A5 G4 G3 U2 C1
Normalized ΔABA
1.4
1.2
1.0
0.8
0.6
0.4
0.2
0.0
(N.D.)

**Figure S6. Insertion and deletion analysis for target #3.**

**(A)** Changes in the ΔABA for all possible insertion (top) and deletion (bottom). For insertions, the line is an average of the four possible insertions at each position. Upper dashed line is the matched target ΔABA and lower dashed line is the RNA-CHAMP detection limit. **(B)** Predicted structure of two insertions at position 3 that relax (3C, top) or further basepair (3U, bottom) with the target RNA. **(C)** Top triangle plot: double deletion analysis. Bottom triangle plot: double insertion analysis.

**A**

Matched target: CCATAGAGAGGTTATCCGCTCA
Encoding sequence: CUATAGAGAGGTTATCCGGTCA

↓

Position 4 substitution-G, Position 21 sustitution-U

↓

| Position | Substitution A | Substitution U | Substitution G | Substitution C | Insertion A | Insertion U | Insertion G | Insertion C | Deletion |
|---|---|---|---|---|---|---|---|---|---|
| 1 | 0 | 0 | 0 | 0 | 0 | 0 | 0 | 0 | 0 |
| 2 | 0 | 0 | 0 | 0 | 0 | 0 | 0 | 0 | 0 |
| 3 | 0 | 0 | 0 | 0 | 0 | 0 | 0 | 0 | 0 |
| 4 | 0 | 0 | 1 | 0 | 0 | 0 | 0 | 0 | 0 |
| ⋮ | | | | | | | | | |
| 21 | 0 | 1 | 0 | 0 | 0 | 0 | 0 | 0 | 0 |
| 22 | 0 | 0 | 0 | 0 | 0 | 0 | 0 | 0 | 0 |

**B**

Encoding sequence: CUGTAGAGAGGTTATCCGCTCA

C21U, A20G

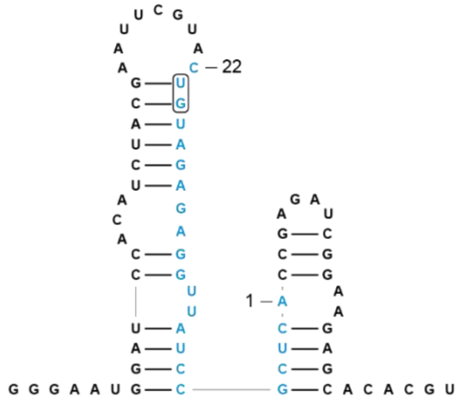


→

| Position | Base pairing |
|---|---|
| 1 | 0 |
| 2 | 1 |
| 3 | 1 |
| 4 | 1 |
| 5 | 1 |
| 6 | 1 |
| 7 | 1 |
| 8 | 1 |
| 9 | 1 |
| 10 | 0 |
| 11 | 0 |
| 12 | 1 |
| 13 | 1 |
| 14 | 0 |
| 15 | 0 |
| 16 | 1 |
| 17 | 1 |
| 18 | 1 |
| 19 | 1 |
| 20 | 1 |
| 21 | 1 |
| 22 | 0 |

**C**

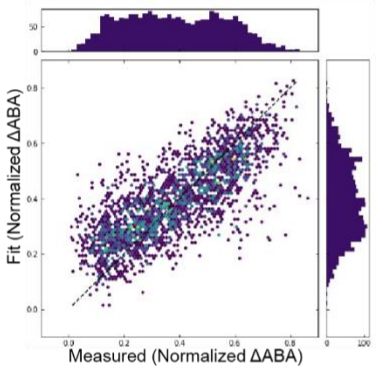


**Figure S7. Cas13d modeling and machine learning model.**
**(A)** Each position along the target has eight possible alterations including substitutions, insertions, and deletions. For example, position four, which is a cytosine in the matched target, can be substituted for A, U or G (C4A, C4U, or C4G), can have one of four insertions (A, U, G, C), or can be deleted. **(B)** Target RNA accessibility is encoded as the number of intramolecularly basepaired nucleotides, as illustrated. RNA structure is predicted by the ViennaRNA package.

**(C)** We trained half of our dataset on a convolutional neural network (CNN) and tested the rest of the dataset with training alteration of 1,000 (epoch). In this machine learning model, there are 37,089 total trained parameters. Pearson's $r = 0.77$.

### SUPPLEMENTARY METHODS

#### Oligonucleotides and DNA libraries

Primers, protospacer flanking sequence DNA libraries, and crRNAs, and target RNAs were purchased from IDT. Mispaired target DNA oligonucleotide libraries were purchased from Twist or GenScript. DNA libraries for probing the protospacer flanking sequence (PFS) were generated via PCR amplification (Q5 High-Fidelity 2X Master Mix, NEB) of mixed based oligo ordered from IDT with primers JK044 and JK045 (**Table S3)**. These mixed based oligoes included three randomized bases on either end of the target RNA. After PCR, Illumina adapters and sequencing primer attachment sites were added for downstream next generation DNA sequencing (NGS). Final amplified libraries were constructed as 5'-P7-SP1-T7 promoter-SP2-TerB-P7. For mispaired DNA oligonucleotide libraries, we designed a custom oligonucleotide DNA pool (purchased from Twist or GenScript). The DNA pool was PCR amplified using primers JK044 and JK045. These primers also added adapters for Illumina-based sequencing. The PFS and mispaired DNA libraries were pooled and sequenced on a conventional MiSeq instrument using a 150-cycle reagent kit v3 (Illumina). To prevent data loss due to sequencing a low diversity library, we also spiked in sheared human cDNA and PhiX DNA to a total of 50% of the sequencing run.

#### Protein Expression and Purification

Eubacterium siraeum Cas13d (EsCas13d) was subcloned into a pET19-based plasmid with an N-terminal 6xHis-TwinStrep-SUMO fusion to generate plasmid the pIF1023 from pET28a-MH6-EsCas13d (Addgene #108303). The nuclease-dead variant (dCas13d) was generated by introducing the following mutations into the HEPN active site: R295A/H300A/R849A/H854A1. The SNAP tag was added at the N-terminus of dCas13d (pIF1024).

Catalytically active and nuclease-dead variants were purified using the same protocol. Briefly, the over expression plasmid was transformed into BL21 star (DE3) cells (Thermo Fisher). Cells were inoculated in LB containing carbenicillin to OD600~0.7 and induced with 200 mM isopropyl β-d-1-thiogalactopyranoside (IPTG) at 18 °C for 18 hours. Cells were then pelleted, resuspended in the lysis buffer (50 mM HEPES pH 7.4, 500 mM NaCl, 1 mM EDTA, 5% glycerol, 0.1% Tween-20, 1 mM DTT, cOmplete-EDTA-free protease inhibitor cocktail

(Sigma Aldrich), 1 mg ml-1 lysozyme, 2.5 U ml-1 DNaseI, 2.5 U ml-1 salt active nuclease), and lysed completely by sonication. Clarified lysate was applied to a Strep-Tactin Superflow gravity column (IBA Life Sciences). The Strep-Tactin resin was washed with 20 column volumes (CVs) of wash buffer (50 mM HEPES pH 7.4, 500 mM NaCl, 5% glycerol, 1 mM DTT), eluted with 5 CVs of elution buffer (50 mM HEPES pH 7.4, 500 mM NaCl, 10% glycerol, 5 mM D-desthiobiotin, 1 mM DTT), and then concentrated by a spin concentrator (Amicon 30 kDa cutoff Ultra 15, Millipore). Concentrated samples were incubated with homemade SUMO protease for tag cleavage, and with SNAP-surface 488 dye (NEB), if used for RNA-CHAMP experiments at 4°C for 20 hours. Samples were then further purified by a size-exclusion column (Superdex 200 Increase 10/300 GL, GE Life Sciences) using SEC buffer (50 mM Tris-HCl pH 7.5, 500 mM NaCl, 10% glycerol, 2 mM DTT).

For ribonuclear protein (RNP) reconstitution, purified dCas13d was incubated with a six-fold excess of CRISPR RNA (crRNA; from IDT) at 37 °C for 1 hour in RNP buffer (50 mM Tris-HCl pH 7.5, 100 mM NaCl, 6 mM $MgCl_2$, 1 mM DTT), and again subjected to size-exclusion column (Superdex 200 Increase 10/300 GL, GE Life Sciences) to further separate the RNP from the crRNA by using RNP SEC buffer (50 mM Tris-HCl pH 7.5, 150 mM NaCl, 1 mM $MgCl_2$, 10% glycerol, 1 mM DTT). RNP fractions were pooled, spin concentrated (Amicon 30 kDa cutoff Ultra 15, Millipore), flash frozen in liquid nitrogen, and stored at -80°C.

**RNA-CHAMP**

MiSeq chips were collected after sequencing and stored at 4˚C in storage buffer (10 mM Tris-HCL pH 8.0, 1 mM EDTA, 500 mM NaCl) until needed. The chips were placed on a custom-designed microscope stage adapter with integrated microfluidics. The buffer perfusion flow rate was controlled via an automated syringe pump (KD scientific) and kept constant at 100 µl $min^{-1}$ for all washing steps. The schematics and CAD files for the microscope stage designs and all additional components are available on https://github.com/finkelsteinlab/RNA-CHAMP.

The chip surface was regenerated after sequencing to remove leftover fluorescent nucleotides and the synthesized strand. The chip was denatured with 500 µl of 0.1 N NaOH and washed with 500 µl TE buffer (10 mM Tris-HCl pH 8.0, 1 mM EDTA). The chip was then incubated with 500 nM regeneration primers IF363 and IF443 (see **Table S3)** in hybridization buffer (5X SSC [750 mM NaCl, 75 mM sodium citrate pH 7.0], 0.1% Tween-20) for 5 minutes at 85 °C, cooled to 65 °C over 10 minutes, cooled to 40 °C over 30 minutes, and held at 40 °C for 10 mins. During the last 10 minutes at 40 °C, the chip was washed with 1 mL wash buffer (0.3X SSC [45 mM NaCl, 4.5mM sodium citrate pH 7.0], 0.1% Tween-20) to remove unannealed primers.

For most Cas13d binding experiments, concentration gradient of 0.125 nM, 0.25 nM, 0.5 nM, 1 nM, 2 nM, 4 nM, 8 nM, 16 nM, 32 nM, 64 nM and 128 nM of Cas13d were sequentially incubated in the chip. At each concentration, Cas13d were incubated for 10 minutes at 25°C. Then Cas13d was washed out by 300 µl of protein buffer (40 mM Tris-HCl pH 7.5, 150 mM NaCl, 6 mM $MgCl_2$, 1 mM DTT, 0.1 % Tween-20, 0.2 mg/ml BSA). The fluorescent imaged was acquired on a TIRF microscopy in a previous reported setup[1]. After every imaging experiment, chips were treated with protease K (80 units/ml) diluted in TE buffer (10 mM Tris-HCl, 500 mM EDTA) for 30 minutes at 42 °C.

**RNA-CHAMP Data Analysis**

Raw images are run through a CHAMP alignment and intensity calculation pipeline[1]. To background subtract the fluorescent buildup on the surface of the chip, the signal from clusters without a T7 promoter was subtracted from the signal for clusters corresponding to RNA library members. Sequences that were represented by five or more physical RNA clusters were globally fit via the Hill equation without cooperativity to calculate the apparent $K_d$, $I_{max}$, and $I_{min}$:

$$I_{obs} = \frac{I_{max} - I_{min}}{1 + \frac{K_d}{x}} + I_{min}$$

Where $I_{min}$ is the minimum intensity for the fit, $I_{max}$ is the maximum intensity for the fit. $x$ is the concentration, and $I_{obs}$ is the observed intensity. To prevent over-interpret the fitting result, sequences that showed maximum fluorescence intensities below 20% of the matched target intensity were considered "weak binders" and not included in our analysis. Only sequences within our detection limit were included in the analysis (e.g., data within the dashed line in **Fig. 1E**). We report these as not determined (N.D.). Apparent $K_d$ was transformed to change in the apparent binding affinity (ΔABA) by $log\left(\frac{K_{d(mt)}}{K_{d(s)}}\right)$, where $K_{d(s)}$ is the Apparent $K_d$ of a library sequence and $K_{d(mt)}$ is the apparent $K_d$ of the matched target. Finally, all experiments were repeated two or more times. For RNA structure prediction, RNA structures were predicted by RNAfold from ViennaRNA[2] using default settings.

**Biolayer Interferometry**

Binding kinetics were assessed via biolayer interferometry on an Octet RED96e (FortéBio). Biotinylated RNA was immobilized on strepavidin biosensors (FortéBio). The biosensors were then immersed in the 100 nM, 50 nM, and 25 nM of dCas13d RNP complex for 600 seconds as the association step and transferred into binding buffer for 600 seconds to measure Cas13d RNP dissociation. We also acquired the signal from a reference sensor without any dCas13d RNP. This trace was treated as a baseline and subtracted from all other association and dissociation curves. The $k_a$, $k_d$, and $K_d$ values were calculated from global fitting to all the binding curves by using Octet data analysis software v11.1. All BLI measurements are summarized in **Table S1**.

**Collateral Cleavage Fluorescent Assay**

Catalytic active Cas13d was purified as described above. 50 nM of Cas13d were incubated with 50 nM of poly-U reporter (5'-6-FAM-UUUUU-Iowa Black® FQ-3', IDT) and 5 nM of the indicated target RNA (IDT). The reaction was incubated in a 96-well plate in the RT-PCR system (ViiA 7) at 25°C. Fluorescent intensities were detected every minute for a total duration of 120 minutes. Technical duplicates were done in every plate, and two or three biological replicates were done for each sequence. Mean of the technical duplicate of one experiment were showed in the plot. Initial slope at 20 minutes time point were calculated to quantitatively compare the cleavage activity. All fluorescent cleavage data are summarized in **Table S2**.

**Computational Modeling**

To extract mechanistic insights into off-target RNA binding, we created generalized models across all target RNA experiments. First, all ΔABAs were normalized to be between 1 and 0 for the upper and lower detection limits, respectively. Model I solely considers basepairing across the 22 nucleotides target RNA according to the function below. The RNA structure was predicted by ViennaRNA[2]. The model adjusts the 22 parameters $a_i$, one for each base in the target RNA sequence:

$$f_{BP}(i) = \begin{cases} 1, & if\ position\ i\ was\ based\ paired\ with\ other\ bases \\ 0, & Otherwise \end{cases}$$

$$\text{Model I: } \widehat{k_{\Delta ABA}} = \sum_{i=1}^{N} a_i * f_{BP}(i)$$

Model II includes an additional term, $g(k)$, the predicted minimal free energy (MFE) in kcal $mol^{-1}$ of sequence $k$ (predicted by RNAfold from ViennaRNA[2]).

$$\text{Model II: } \widehat{k_{\Delta ABA}} = \sum_{i=1}^{N} a_i * f_{BP}(i) + e * g(k)$$

Model III is the relative encoding-only model and has three main terms that summarize the relative penalties for insertions (*I*), deletions (*D*), or mismatches (*M*) in the target sequence relative to the matched target. As an example, consider a sequence with C2G and U10A alteration compared to the matched target strand, $k_{mt}$. These operations can be conceptually written as:

$$\{(Mismatch, 2, \mathrm{G}), (Mismatch, 10, A)\}$$

Thus $f_M(2, G)$ and $f_M(10, A)$ would evaluate to 1 and all other inputs for $f_x$ would evaluate to 0. There was a total of 9 parameters for each RNA positions: deletion, insertion A, insertion U, insertion G, insertion C, mismatch A, mismatch U, mismatch G, and mismatch C. Here, *i* denotes the sequence position, *v* is the altered RNA base identity, and *b, c, d* are the three sets of adjustable parameters.

$$f_x(i, v) = \begin{cases} 1, & if\ oper\ x\ \ used\ to\ transfrom\ k_{mt}\ to\ k \\ 0, & Otherwise \end{cases}$$

$$\text{Model III: } \widehat{k_{\Delta ABA}} = \sum_{i\,\in\,I} b_{i,v} * f_I(i, v) + \sum_{i\,\in\,D} c_i * f_D(i, 0) + \sum_{i\,\in\,M} d_{i,v} * f_M(i, v)$$

Model IV includes an additional MFE term, $g(k)$, as previously described.

$$\text{Model IV: } \widehat{k_{\Delta ABA}} = \sum_{i\,\in\,I} b_{i,v} * f_I(i, v) + \sum_{i\,\in\,D} c_i * f_D(i, 0) + \sum_{i\,\in\,M} d_{i,v} * f_M(i, v) + e * g(k)$$

Model V is the combination of Model I and Model III that includes both target RNA basepairing and relative mismatch/indel encoding.

$$\text{Model V: } \widehat{k_{\Delta ABA}} = \sum_{i=1}^{22} a_i * f_{BP}(i) + \sum_{i\,\in\, I} b_{i,v} * f_I(i,v) + \sum_{i\,\in\, D} c_i * f_D(i,0) + \sum_{i\,\in\, M} d_{i,v} * f_M(i,v)$$

Model VI has the following parameters: base pairing, relative encoding, and the MFE of the predicted lowest-energy structure. The normalized ΔABA for partially matched RNAs, $k$, that are related to the matched target, $k_{mt}$, were modeled using a linear combination of the following features. $f_x(i,v)$ denotes different types of sequence alteration and RNA accessibility in the model, where *BP, I, D,* and *M* were base pairing, insertions, deletions, and mismatches respectively.

$$\text{Model VI: } \widehat{k_{\Delta ABA}} = \sum_{i=1}^{22} a_i * f_{BP}(i) + \sum_{i\,\in\, I} b_{i,v} * f_I(i,v) + \sum_{i\,\in\, D} c_i * f_D(i,0) + \sum_{i\,\in\, M} d_{i,v} * f_M(i,v) + e * g(k)$$

The weights of the terms $a_i$, $b_{i,v}$, $c_i$, $d_{i,v}$, and *e* are the adjustable parameters that are used to fit the experimental data and represent the penalties of each operational transformation on altered sequences *k* in the library.

Ridge regression was used to determine the weights of our parameters to fit to the experimental training set. Ridge regression is a variant of linear regression which attempts to minimize the training loss value of the expression:

$$\sum_{k\,\in T_{tr}} (M(k) - k_{ABA})^2 \ + \lambda \sum_{\beta\,\in\, X} \beta^2$$

Where M is defined to be the model for predicting ABA with all the weights $\beta$ being values in set X = $\{a_i, b_i, c_i, d_i, e\}$. The predicted values from Model M are compared to the measured ΔABAs values, $k_{\Delta ABA}$: the smaller the absolute difference between the two the greater model's accuracy. Ridge regression helps maintain robustness of linear models and prevents overfitting by penalizing arbitrarily large weights. The parameter $\lambda$ at which the weight values in the model appear to stabilize is around 1 which was used throughout all models.

Finally, we considered a sequential convoluted neuron network (CNN) model. The model was built on a single Conv2D layer with 64 filters and the kernel size of 54x1[3]. Then, a final layer of MaxPooling2D with a pool size of 3x1 was added. The CNN model was trained by the same dataset used in the simple linear model. The dataset has 4862 sequences, and only half of the sequences were used for training the model. The model was trained through 1000 epochs and were tested on the rest of the sequences.

### Supplemental Tables

### Table S1. Biolayer interferometry and RNA cleavage rates for select RNAs

| **Sequences** | $K_d$ **(M)** | $K_d$ **Error** | $k_a$ **($M^{-1}S^{-1}$)** | $k_a$ **Error** | $k_d$ **($S^{-1}$)** | $k_d$ **Error** |
|---|---|---|---|---|---|---|
| MT | 3.1E-09 | 1.8E-11 | 1.7E+05 | 6.4E+02 | 5.2E-04 | 2.5E-06 |
| C2G | 2.0E-09 | 1.6E-11 | 2.0E+05 | 8.1E+02 | 4.0E-04 | 2.6E-06 |
| C4A | 1.6E-09 | 1.0E-11 | 2.2E+05 | 6.6E+02 | 3.5E-04 | 1.9E-06 |
| G5U | 9.3E-09 | 1.2E-10 | 4.5E+04 | 3.5E+02 | 4.2E-04 | 4.1E-06 |
| C7A | 2.1E-09 | 1.3E-11 | 2.2E+05 | 7.9E+02 | 4.5E-04 | 2.3E-06 |
| U10G | 3.8E-09 | 2.9E-11 | 1.3E+05 | 6.3E+02 | 5.1E-04 | 3.1E-06 |
| G15A | 3.4E-09 | 1.9E-11 | 1.8E+05 | 6.4E+02 | 6.0E-04 | 2.4E-06 |
| A18G | 8.7E-09 | 4.8E-11 | 7.3E+04 | 2.8E+02 | 6.3E-04 | 2.5E-06 |
| A20C | 6.3E-09 | 3.4E-11 | 10.0E+04 | 3.7E+02 | 6.3E-04 | 2.5E-06 |
| C22A | 3.6E-09 | 2.0E-11 | 1.9E+05 | 7.2E+02 | 6.6E-04 | 2.6E-06 |
| 5'-UAA | 6.0E-09 | 5.4E-11 | 1.0E+05 | 6.2E+02 | 6.1E-04 | 4.0E-06 |
| 3'-GCU | 4.2E-09 | 2.0E-11 | 1.7E+05 | 5.9E+02 | 7.2E-04 | 2.3E-06 |
| 3'-GGU | 7.1E-09 | 5.8E-11 | 1.2E+05 | 7.3E+02 | 8.4E-04 | 4.3E-06 |
| 3'-UAA | 6.0E-09 | 5.4E-11 | 1.0E+05 | 6.2E+02 | 6.1E-04 | 4.0E-06 |
| 3'-UGG | 8.6E-09 | 1.1E-10 | 7.0E+04 | 6.0E+02 | 6.0E-04 | 5.4E-06 |
| 3'-GUU | 4.4E-09 | 3.5E-11 | 2.5E+05 | 1.6E+03 | 1.1E-03 | 4.7E-06 |

### Table S2. Slope of fluorescent cleavage assay for selected RNAs

| **Sequences** | **Slope (AU/min)** | **S.D.** |
|---|---|---|
| MT | 22000 | 190 |
| C2G | 580 | 110 |
| C4A | 590 | 26 |
| G5U | 900 | 180 |
| C7A | 3300 | 430 |
| U10G | 8300 | 200 |
| G15A | 16000 | 810 |
| A18G | 12000 | 1800 |
| A20C | 15000 | 2000 |
| C22A | 13000 | 1200 |

### Table S3. DNA oligonucleotides used in this study

| Name | Type | Description | Sequence (5’-3’) |
|---|---|---|---|
| Library extension primer-Forward | Oligo | Extend library oligo with illumine adapters | AATGATACGGCGACCACCGAGATCTACACTCTTTCCCTACACGACGCT CTTCCGATCT |
| Library extension primer-Reverse | Oligo | Extend library oligo with illumine adapters | CAAGCAGAAGACGGCATACGAGATGAACAACATGACGTGACTTTAGT TACAACATACTAATTGTGACTGGAGTTCAGACGTGTGCTCTTCCGATCT |
| Target 1 | Oligo pool | 6N PFS oligo library | CCTACACGACGCTCTTCCGATCTTAATACGACTCACTATAGGGAATGG ATCCACATCTACGAATTCNNNCCATAGAGAGGTTATCCGCTCANNNAG ATCGGAAGAGCACACGTCTGAAC |
| Target 2 | Oligo pool | 6N PFS oligo library | CCTACACGACGCTCTTCCGATCTTAATACGACTCACTATAGGGAATGG ATCCACATCTACGAATTCNNNGTTGTTCTCCGTCTATAAATACNNNAG ATCGGAAGAGCACACGTCTGAAC |
| t7_promoter_scramble | Oligo | T7 promoter scramble oligo | CCTACACGACGCTCTTCCGATCTACGGTAGATCTAAAGTCACTAATGG ATCCACATCTACGAATTCNNNTTTGATCTGAAATATTCAGGTCNNNAG ATCGGAAGAGCACACGTCTGAAC |
| Non-target negative control | Oligo | Non-target oligo as a negative control | CCTACACGACGCTCTTCCGATCTTAATACGACTCACTATAGGGAATGG ATCCACATCTACGAATTCGTTAGCTAGAAGGGGAAGTTGGTTATGGAG ATCGGAAGAGCACACGTCTGAAC |
| P7 regeneration primer (IF363) | Oligo | Regenerate all cluster into dsDNA on the chip | CAAGCAGAAGACGGCATACGAGAT |
| PhiX labeling primer (IF443) | 5’ Digoxigenin labeled Oligo | Regenerate PhiX clusters on the chip | /5DigN/CGGTCTCGGCATTCCTGCTGAACCGCTCTTCCGATC |

### Table S4. RNA oligonucleotides used in this study

| Name | Type | Description | Sequence (5'-3') |
|---|---|---|---|
| crRNA_ Target 1 | crRNA | PFS analysis (Fig. 2), Mismatch analysis (Fig. 3) | CACCCGUGCAAAAAUGCAGGGGUCUAAAACUGAGCGGAUAAC CUCUCUAUGG |
| crRNA_ Target 2 | crRNA | PFS analysis (Fig. S2) | CACCCGUGCAAAAUUGCAGGGGUCUAAAACCUUCUCCAAAUU GUUUCAUCCU |
| crRNA_ Target 3 | crRNA | Mismatch analysis (Fig. S3) | CACCCGUGCAAAAUUGCAGGGGUCUAAAACGACCUGAAUAUU UCAGAUCAAA |
| MT | 3' biotinylated ssRNA | BLI, Fluorescent cleavage assay | CCACAUCUACGAAUUCGUACCAUAGAGAGGUUAUCCGCUCAC CGAGAUCGGAAGAGCACA/3Bio/ |
| C2G | 3' biotinylated ssRNA | BLI, Fluorescent cleavage assay | CCACAUCUACGAAUUCGUACCAUAGAGAGGUUAUCCGCUGAC CGAGAUCGGAAGAGCACA/3Bio/ |
| C4A | 3' biotinylated ssRNA | BLI, Fluorescent cleavage assay | CCACAUCUACGAAUUCGUACCAUAGAGAGGUUAUCCGAUCAC CGAGAUCGGAAGAGCACA/3Bio/ |
| G5U | 3' biotinylated ssRNA | BLI, Fluorescent cleavage assay | CCACAUCUACGAAUUCGUACCAUAGAGAGGUUAUCCUCUCAC CGAGAUCGGAAGAGCACA/3Bio/ |
| C7A | 3' biotinylated ssRNA | BLI, Fluorescent cleavage assay | CCACAUCUACGAAUUCGUACCAUAGAGAGGUUAUACGCUCAC CGAGAUCGGAAGAGCACA/3Bio/ |
| U10G | 3' biotinylated ssRNA | BLI, Fluorescent cleavage assay | CCACAUCUACGAAUUCGUACCAUAGAGAGGUGAUCCGCUCAC CGAGAUCGGAAGAGCACA/3Bio/ |
| G15A | 3' biotinylated ssRNA | BLI, Fluorescent cleavage assay | CCACAUCUACGAAUUCGUACCAUAGAAAGGUUAUCCGCUCAC CGAGAUCGGAAGAGCACA/3Bio/ |
| A18G | 3' biotinylated ssRNA | BLI, Fluorescent cleavage assay | CCACAUCUACGAAUUCGUACCAUGGAGAGGUUAUCCGCUCAC CGAGAUCGGAAGAGCACA/3Bio/ |
| A20C | 3' biotinylated ssRNA | BLI, Fluorescent cleavage assay | CCACAUCUACGAAUUCGUACCCUAGAGAGGUUAUCCGCUCACC GAGAUCGGAAGAGCACA/3Bio/ |
| C22A | 3' biotinylated ssRNA | BLI, Fluorescent cleavage assay | CCACAUCUACGAAUUCGUAACAUAGAGAGGUUAUCCGCUCAC CGAGAUCGGAAGAGCACA/3Bio/ |

| | | | |
|---|---|---|---|
| 5’-UAA | 3’ biotinylated ssRNA | BLI | CCACAUCUACGAAUUCUAACCAUAGAGAGGUUAUCCGCUCACCGAGAUCGGAAGAGCACA/3Bio/ |
| 3’-GCU | 3’ biotinylated ssRNA | BLI | CCACAUCUACGAAUUCGUACCAUAGAGAGGUUAUCCGCUCAGCUAGAUCGGAAGAGCACA/3Bio/ |
| 3’-GGU | 3’ biotinylated ssRNA | BLI | CCACAUCUACGAAUUCGUACCAUAGAGAGGUUAUCCGCUCAGGUAGAUCGGAAGAGCACA/3Bio/ |
| 3’-UAA | 3’ biotinylated ssRNA | BLI | CCACAUCUACGAAUUCGUACCAUAGAGAGGUUAUCCGCUCAUAAAGAUCGGAAGAGCACA/3Bio/ |
| 3’-UGG | 3’ biotinylated ssRNA | BLI | CCACAUCUACGAAUUCGUACCAUAGAGAGGUUAUCCGCUCAUGGAGAUCGGAAGAGCACA/3Bio/ |
| 3’-GUU | 3’ biotinylated ssRNA | BLI | CCACAUCUACGAAUUCGUACCAUAGAGAGGUUAUCCGCUCAGUUAGAUCGGAAGAGCACA/3Bio/ |
| Poly-U reporter | 5’ Fluorescein, 3’ quencher ssRNA | Fluorescent reporter for the cleavage assay | 6-FAM-UUUUU-Iowa Black® FQ |
